# WHIRLY1 of barley and maize share a PRAPP motif conferring nucleoid compaction

**DOI:** 10.1101/2020.10.19.344945

**Authors:** Svenja Oetke, Axel J. Scheidig, Karin Krupinska

## Abstract

WHIRLY1 in barley was shown to be a major architect of plastid nucleoids. Its accumulation in cells of *E. coli* coincided with an induction of nucleoid compaction and growth retardation. While WHIRLY1 of maize had similar effects on *E. coli* cells, WHIRLY1 proteins of Arabidopsis and potato a well as WHIRLY2 proteins had no impact on nucleoid compaction in *E. coli*. By mutagenesis of *HvWHIRLY1* the PRAPP motif at the N-terminus preceding the highly conserved WHIRLY domain was identified to be responsible for the nucleoid compacting activity of HvWHIRLY1 in bacteria. This motif is found in WHIRLY1 proteins of most members of the Poaceae family, but neither in the WHIRLY2 proteins of the family nor in any WHIRLY protein of eudicot species such as *Arabidopsis thaliana*. This finding indicates that a subset of the monocot WHIRLY1 proteins has acquired a specific function as nucleoid compacters by sequence variation in the N-terminal part preceding the conserved WHIRLY domain and that in different groups of higher plants the compaction of nucleoids is mediated by other proteins.

## INTRODUCTION

WHIRLY1 has been identified as a major protein of chloroplast nucleoids (1–3). Chloroplasts are descendants of cyanobacteria and retained a residual genome with prokaryotic features. In higher plants the plastid genome encodes, however, only a few tens of the several thousand proteins of the organelle (4,5) due to a massive transfer of genes to the nucleus (4,6). Multiple copies of the small organelle genome are organized in structures resembling bacterial nucleoids (7,8). In contrast to bacteria, plastids have multiple nucleoids, which are known to change in number and morphology during development (7,9,10). Nucleoids in bacteria are organized by several abundant architectural proteins called NAPs (nucleoid associated proteins) such as HU (heat-unstable) or H-NS (histone-like nucleoid structuring) (11,12). While dinoflagellates and rhodophytes still possess bacterial NAPs (13,14), plastid nucleoids of higher plants lack these proteins (8,13,15). Instead, they have acquired eukaryotic proteins (16) that were shown to influence the shaping and organization of plastid nucleoids (10), such as WHIRLY, SWIB and SIR (sulfite reductase) (16). Besides numerous proteins playing roles in DNA/RNA associated processes, nucleoids contain additional proteins that might link DNA/RNA associated functions on one hand and energy metabolism and biosynthetic activities on the other hand (17). Several of the nucleoid associated proteins have a second localization in the nucleus (18). Thereby they are excellent candidates for genome communication involving coordinate changes in nucleoid architecture and chromatin organization. In this respect it is interesting that selected stress related changes in histone modification are suppressed in barley plants with a knockdown of HvWHIRLY1 (19).

WHIRLY proteins are plant specific DNA binding proteins sharing the highly conserved WHIRLY domain. They were shown to form tetramers (20) and to further assemble into hexamers of the tetramers, i.e. 24-mers (21). Most plants have two WHIRLY proteins, whereby WHIRLY1 is targeted to chloroplasts and WHIRLY2 is targeted to mitochondria (22). Some plants such as *Arabidopsis thaliana* (*A. thaliana*) have a third WHIRLY protein which is targeted to chloroplasts (22). While the WHIRLY domain is highly conserved in all plants, the N-terminal and C-terminal parts of the WHIRLY proteins are highly variable suggesting that the proteins are multifunctional (23).

An intriguing feature of WHIRLY1 is its dual localization in plastids and nucleus (24). In both compartments it has functions associated with DNA and/or RNA. In chloroplasts of maize WHIRLY1 was shown to associate to DNA in a sequence non-specific manner and also to bind to a subset of RNAs deriving from intron containing genes (25). In barley, WHIRLY1 was shown to bind to a similar subset of RNAs (26) and to induce compaction of nucleoids (27). In the nucleus it has been identified as a transcriptional activator of the pathogen response genes *PR-10* of potato (28) and *PR-1* of Arabidopsis (29). It has been also shown to influence telomer maintenance (30) and biosynthesis of microRNAs (31). Plants with reduced abundance of WHIRLY1 show a delay of chloroplast development (32) and leaf senescence (33) as well as deficiencies in the responses towards nutrition (34) and light (31). Taken together the data suggest a major role of WHIRLY1 proteins in development and stress resistance.

Investigations with a double mutant for the two plastid WHIRLY proteins of Arabidopsis (*why1 why3*) revealed that in plastids WHIRLY proteins contribute to plastid genome stability by protecting DNA against illegitimate repeat-mediated recombination (35). As a consequence of ptDNA rearrangements, leaves occasionally showed variegation indicative of a disturbance of chloroplast development (36). It has been suggested that WHIRLY proteins resemble eubacterial single stranded DNA binding (SSB) proteins (37) affecting diverse single stranded DNA associated processes such as replication, recombination and repair (38). Furthermore, they were proposed to function as organizational scaffolds recruiting genome maintenance complexes (39). In transgenic barley plants with a RNAi mediated knockdown of *HvWHIRLY1*, chloroplasts contained loosely packed nucleoids which were much larger than in control plants (27). They also showed a higher expression of the plant organelle targeted DNA polymerase (POP) coinciding with more plastid DNA (27). It is likely that the correct architecture of nucleoids is important for the coordinate progression of replication as well as transcription and translation (40). In accordance with the function of WHIRLY1 as a major architect of plastid nucleoids known to serve as platform for ribosomes (40), *why1* mutants of maize were shown to be impaired in assembly of plastid ribosomes. Also the barley plants with a knockdown of *HvWHIRLY1* were reported to have reduced levels of rRNAs (41) being a measure of ribosome abundance in chloroplasts (42).

WHIRLY1 of barley was shown to induce nucleoid compaction both in chloroplasts and in *E. coli* (27). In this study, it has been shown that the compacting impact of WHIRLY1 is specific for members of the Poaceae family. By mutagenesis of the N-terminal part of HvWHIRLY1, a proline-rich motif, i.e. PRAPP, was identified to be responsible for nucleoid compaction. Neither WHIRLY1 proteins of eudicots nor any WHIRLY2 protein contain a PRAPP motif coinciding with a lack of the ability to compact nucleoids. Based on these results it is proposed that nucleoids in different groups of plants are compacted by different proteins.

## MATERIALS AND METHODS

### Bacterial strains and growth conditions

*Escherichia coli* (*E. coli*) DH5α cells were grown in LB broth (43) at 37°C with shaking at 180 rpm (Labshaker, InnovaTM4400, New Brunswick Scientific, Edison, New Jersey, USA). For selection of transformants, the LB medium was supplemented with 100 μg/ml ampicillin. The *Agrobacterium tumefaciens* (*A. tumefaciens*) strain GV3101 (pMP90) was used for stable transformation of *A. thaliana*, ecotype Columbia (44). *A. tumefaciens* was grown for two days at 28°C and 140 rpm in YEB broth (5 g/l yeast extract, 1 g/l beef extract, 5 g/l peptone, 5 g/l sucrose, 2 mM MgSO_4_). Antibiotics were added at a concentration of 100 μg/ml rifampicin, 50 μg/ml gentamicin and 50 μg/ml spectinomycin for selection of strains and plasmids.

### Heterologous expression of *WHIRLY* genes in *E. coli* cells

For heterologous expression, the sequences encoding mature WHIRLY proteins of different species were amplified by polymerase chain reaction using the *Pfu* DNA polymerase (Thermo Fisher Scientific, Waltham, MA, USA) and specific primers (Supplementary Table S1) and were afterwards cloned into the pASK-IBA3 vector via the BsaI restriction site (IBA Life Science, MO, USA): *WHIRLY1* (*HvWHIRLY1*, AK365452.1) and *WHIRLY2* (*HvWHIRLY2*, BF627441.2) from barley (*Hordeum vulgare*), *WHIRLY1* (*ZmWHIRLY1*, Zm.88681) and *WHIRLY2* (*ZmWHIRLY2*, NM_001159117.2) from maize (*Zea mays*), *WHIRLY1* (*AtWHIRLY1*, At1g14410), *WHIRLY2* (*AtWHIRLY2*, At1g71260.1) and *WHIRLY3* (*AtWHIRLY3*, At2g02740) from *A. thaliana*, *WHIRLY1* (*StWHIRLY1*, NM_001288226.1) and *WHIRLY2* (*StWHIRLY2*, NM_001288464.1) from potato (*Solanum tuberosum*). For induction of expression, at an OD_600_ 0.6-0.7 anhydrotetracycline was added to a final concentration of 200 μg/l. The growth rates were measured photometrically at 600 nm every hour (Shimadzu, UV-2501PC, Spectrophotometer, Duisburg, Germany).

### DNA condensation assays with cells of *E. coli*

Staining of nucleoids of *E. coli* cells with 4′,6-diamidino-2-phenylindole (DAPI) was performed as described (2). Cells were observed by fluorescence microscopy with a Zeiss Axiophot microscope (Carl Zeiss, Oberkochen, Germany) using a 100 × oil objective (Plan-APOCHROMAT 100x/1,4 Oil Ph3, Zeiss, Feldbach, Switzerland) and the Zeiss filter set 02 (excitation: G365, beam splitter FT: 395, emission: 420 LP). Images were taken with an Olympus DP71 camera and processed by the *cell^F* software (Olympus Europe).

### Preparation of cell lysates from *E. coli*

For immunological analyses of recombinant proteins, *E. coli* cells collected from 1 ml cell culture were resuspended in 1x PSB (50 mM Tris-HCl, pH 6.8, 50 mM DTT, 10% (v/v) glycerol, 2% (w/v) SDS, 0.1% (w/v) bromophenol blue) and were diluted to the same optical density, i.e. OD_600_ 0.7. The cells were ruptured by heating at 95°C for ten minutes and thereafter chilled on ice. Equal amounts of cell lysates were loaded onto a SDS containing polyacrylamide gel.

### Site-directed mutagenesis of *WHIRLY* genes for heterologous expression

To introduce site-directed substitutions, deletions or insertions in *WHIRLY* coding sequences the QuikChange Lightning Site-Directed Mutagenesis kit (Agilent Technologies, Santa Clara, USA) was used according to the manufacturer’s instructions with minor modifications. Annealing was done at 60°C for ten seconds and the cycle number was eighteen. Mutagenic primers were designed based on the QuikChange Primer Design Program (www.agilent.com/genomics/qcpd). All primer sequences that were used for site-directed mutagenesis are listed in Supplementary Table S2.

### Determination of recombinant protein solubility

Two hours after induction of expression 100 ml cultures were harvested by centrifugation for 5 minutes at 12.000 × g and 4°C. The cells were resuspended in 1 ml 1x PBS (137 mM NaCl, 2.7 mM KCl, 10 mM Na_2_HPO_4_ × 12 H_2_O, 2 mM KH_2_PO_4_) per 0.1 g wet weight and sonicated six times 20 seconds at 70% intensity (Bandelin SONOPLUS, Berlin, Germany) with 60 seconds pauses on ice. To separate the soluble (supernatant) from the insoluble fraction (pellet), the lysate was centrifuged for 20 minutes at 11.000 rpm and 4°C. The supernatant was collected in a new tube. The pellet was resuspended in the same volume 1x PBS as used for resuspension of the cells.

### RNA purification from *E. coli*

Two hours +/− induction with anhydrotetracycline 10^10^ cells were transferred to a reaction tube. Per 1 ml culture 200 μl stop mix (5 % phenol in 100 % ethanol) was added. The sample was centrifuged for 5 minutes at 14.000 × g and 4°C. The pellet was resuspended on ice in 1 ml NucleoZol (Macherey-Nagel, Düren, Germany) and incubated for 10 minutes at 65°C with shaking at 250 rpm. 400 μl chloroform/isoamylalkohol (24:1) was added and the sample was mixed by inverting for 10 seconds. Afterwards the sample was centrifuged for 10 minutes at 14.000 × g and 4°C. The aqueous phase was transferred to a new reaction tube and 450 μl phenol/chloroform/isoamylalkohol (25:24:1) were added. The sample was mixed by inverting for 10 seconds and centrifuged for 10 minutes at 14.000 × g and 4°C. For RNA precipitation, the aqueous phase was transferred to a new reaction tube and 1 volume ice-cold isopropanol and 20 μl 3 M Na-Acetat pH 5,2 were added and mixed by inverting. The RNA was incubated at −20°C over night. The next day the RNA was pelleted by centrifugation for 30 minutes at 14.000 × g and 4°C. The RNA pellet was washed twice with 350 μl ice-cold 75 % ethanol and centrifugation for 5 minutes at 14.000 × g and 4°C. The pellet was dryed for 5-10 min at room temperature and resuspended in 30 μl sterile water.

### Determination of mRNA levels in *E. coli* by qRT-PCR

Isolated total RNA from *E. coli* was used for cDNA synthesis employing QuantiTect^®^ Reverse Transcriptase Kit (Qiagen, Hilden, Germany) according to the manufacturer’s protocol. Quantitative real time PCR (qRT-PCR) analyses were performed with the QuantiFast SYBR Green PCR Kit (Qiagen, Hilden, Germany) according to the manufacturer’s protocol using gene specific primers (Supplementary Table S3). Data analysis was accomplished by the Rotor-Gene Q software version 2.0.2.4 (Qiagen, Hilden, Germany). Relative quantification of transcript levels was performed using the “Delta-delta CT method” as presented by PE Applied Biosystems (Perkin Elmer, Foster City, CA, USA). Data were normalized to the *cysG* transcript and data are shown in relation to *E. coli* cells containing the empty vector pASK-IBA3 without induction.

### Plant material

Seeds of *A. thaliana*, ecotype Columbia, were surface sterilized according to (45) and afterwards sown on MS agar plates (4.4 g/l Murashige & Skoog med. incl. Mod. Vitamins M0245 (Duchefa Haarlem, The Netherlands), 3% (w/v) sucrose, 0.75% (w/v) phyto agar, pH 5.8) in sterile boxes (ECO2box / green filter (OVAL model 80mm H), Duchefa, Haarlem, The Netherlands). Seedlings were cultivated in a climate cabinet at 21°C with 16 h illumination (60-80 μmol photons s^−1^ m^−2^). At 14 days after sowing, seedlings were either used for microscopic analyses or were transferred in Jiffypots^®^ (Jiffy Products International AS, Norway) and grown for additional two weeks in a climate chamber at 21°C with 16 h illumination (50-100 μmol photons s^−1^ m^−2^). For protein extraction, rosette leaves of 4 week-old plants were used. In case of stable transformation with *A. tumefaciens*, plants were grown for 5-6 weeks on Jiffypots in a climate chamber at 21°C with 16 h illumination (50-100 μmol photons s^−1^ m^−2^).

### Generation of AtWHIRLY1+PRAPP:HA overexpressing lines

To generate transgenic lines overexpressing AtWHIRLY1+PRAPP:HA, the coding sequence was cloned in the entry vector pENTR/D-TOPO (Thermo Fisher Scientific, Waltham, MA, USA). Afterwards the modified WHIRLY sequence was transferred by LR reaction into the binary vector pB2GW7,0 (Department of Plant Systems Biology (PSB), University of Gent, Belgium) according to manufacturer’s instructions (Thermo Fisher Scientific, Waltham, MA, USA). The resulting plasmid was introduced into competent *A. tumefaciens* cells by electroporation. For electroporation cells were thawed on ice, 10 ng plasmid were added and carefully mixed. Cells were transferred to pre-chilled cuvettes. For electroporation the following settings were chosen: 2.5 kV, 25 μF, 400 Ω (Gene Pulser, Bio-Rad Laboratories, Munich, Germany). One ml YEB medium was added after electroporation and the cells were transferred on ice. Then the cells were incubated for three to four hours at 28°C and 140 rpm and afterwards plated on selective YEB plates (100 μg/ml rifampicin, 50 μg/ml gentamicin and 50 μg/ml spectinomycin).

Stable transformation of *A. thaliana* (ecotype Columbia) was performed according to the protocol of (46) with little modifications. After overnight growth at 28°C and 140 rpm, *A. tumefaciens* cells were collected by centrifugation for five minutes at 2.800 × *g* and 4°C. Cells were resuspended in inoculation medium (5% (w/v) sucrose and 0.05% (v/v) Silwet-L 77) and diluted to a final inoculation concentration of OD_600_ of 0.5. Inoculation of Arabidopsis was performed by pipetting the bacterial suspension on buds and immature flowers. This procedure was repeated twice at 7-day intervals. Selection of transformed progeny seedlings was done by treatment with the herbicide glufosinate (Basta^®^ was diluted 1:1,000 with tap water).

### Extraction of proteins and isolation of plastids from plant material

Proteins were extracted from seedlings with a buffer consisting of 62.5 mM Tris, pH 6.8, 10% (v/v) glycerol, 1% (w/v) SDS and 5% (v/v) ß-mercaptoethanol. Plastids were isolated from four week old plants using percoll gradients as described (47). Protein concentrations were determined using the Bio-Rad Assay according to the manufacturer’s instructions (Bio-Rad Laboratories, Munich, Germany) based on the method of (48).

### Immunoblot analysis

Equal amounts of proteins or bacterial cell lysates were subjected to SDS-PAGE on 14% (w/v) polyacrylamide gels according to the protocol of (49). Separated proteins were stained with colloidal Coomassie Brilliant Blue (CBB) (50). Proteins were blotted onto nitrocellulose by semi-dry electroblotting and treated as described (51). To check the transfer of the proteins, the membrane was treated with Ponceau S (0.2% (w/v) (Roth, Karlsruhe, Germany), 3% (w/v) trichloroacetic acid) and successively washed in distilled water until the protein bands were clearly visible. Primary antibodies used in this study were directed toward barley WHIRLY1 (24), maize WHIRLY1 (provided by J. Pfalz, University of Jena, Germany), the HA tag (Roche, Mannheim, Germany) and LHCB5 (No. 0202235431, Agrisera, Vännäs, Sweden). For immunological detection of recombinant proteins from Arabidopsis and potato the Strep-Tactin®-HRP conjugate (IBA, Göttingen, Germany) was used. Immunoreactions were detected by chemoluminescence using different kits (GE Healthcare, Buckinghamshire, UK; Thermo Scientific, Waltham, MA, USA; Lumigen, Southfield, MI, USA).

### Staining of nucleoids with SYBR Green I

For staining of nucleoids with SYBR Green I (SYBR™ Green I Nucleic Acid Gel Stain, 10,000X concentrate in DMSO, Invitrogen, Karlsruhe, Germany), cross-sections from the first true leaves of 14 days old Arabidopsis seedlings were excised and fixed overnight at 4°C in 4% (w/v) formaldehyde (prepared from paraformaldehyde). The sections were washed three times with 2x SSC (0.3 M NaCl, 30 mM sodium citrate, pH 7.0) and stained with SYBR Green I in 2x SSC in a concentration of 1:1,000 for 30 minutes at room temperature in the dark. After washing with 2x SSC for 30 minutes at room temperature in the dark, the cross-sections were embedded in a solution consisting of 50% (v/v) glycerol and 1x SSC. Microscopy was performed with a confocal laser-scanning microscope (Leica TCS SP5, Leica Microsystems, Wetzlar, Germany), employing a 63x 1.2 water objective HCX PLAPO and the LAS AF –Software. Fluorescence was excited at 488 nm (15% intensity) using an argon laser. The fluorescence of SYBR Green I was detected at emissions of 508-536 nm. Maxiprojections were taken with a pinhole airy of 1, the size-depth was 4.0 μm with 4 steps.

### Sequence analysis

Multiple sequence alignments were performed with Geneious 8.1.6 using Clustal W.

## RESULTS

### Overexpression of the *WHIRLY1* genes of barley and maize in *E. coli* coincides with compaction of bacterial nucleoids

Overexpression of the barley *WHIRLY1* sequence encoding the mature protein without the plastid target peptide (PTP) was previously shown to induce compaction of bacterial nucleoids coinciding with reduced growth of the bacterial cells (27). To test whether maize WHIRLY1 (ZmWHIRLY1) does also induce compaction of nucleoids, the ZmWHIRLY1 sequence without PTP was overexpressed in *E. coli*. Staining of the cells with DAPI revealed that nucleoids in cells overexpressing the *ZmWHIRLY1* gene contain condensed nucleoids resembling those in the cells accumulating the recombinant HvWHIRLY1 protein (Figure 1A). Compaction coincided with a reduced growth of cells as reported earlier for HvWHIRLY1 (Figure 1B) (27). To investigate whether the recombinant protein is produced by the bacteria, specific antibodies directed against ZmWHIRLY1 (Figure 1C) and HvWHIRLY1 (Supplementary Figure S1), respectively, were used. ZmWHIRLY1 was found to be detectable already one hour after induction of protein synthesis (Figure 1C). The antibody directed against ZmWHIRLY1 (25) recognized additionally a second protein of slightly higher molecular weight which is already present in cells before induction of protein synthesis and hence cannot be related to ZmWHIRLY1. In case of barley the recombinant protein of about 27 kDa was most prominent already after one hour of induction. In addition, a second protein with slightly higher molecular weight was detected (Supplementary Figure S1). This could be a post-translationally modified form of HvWHIRLY1. During further growth of the cells low molecular weight proteins likely representing degradation products accumulated.

**Figure 1.**
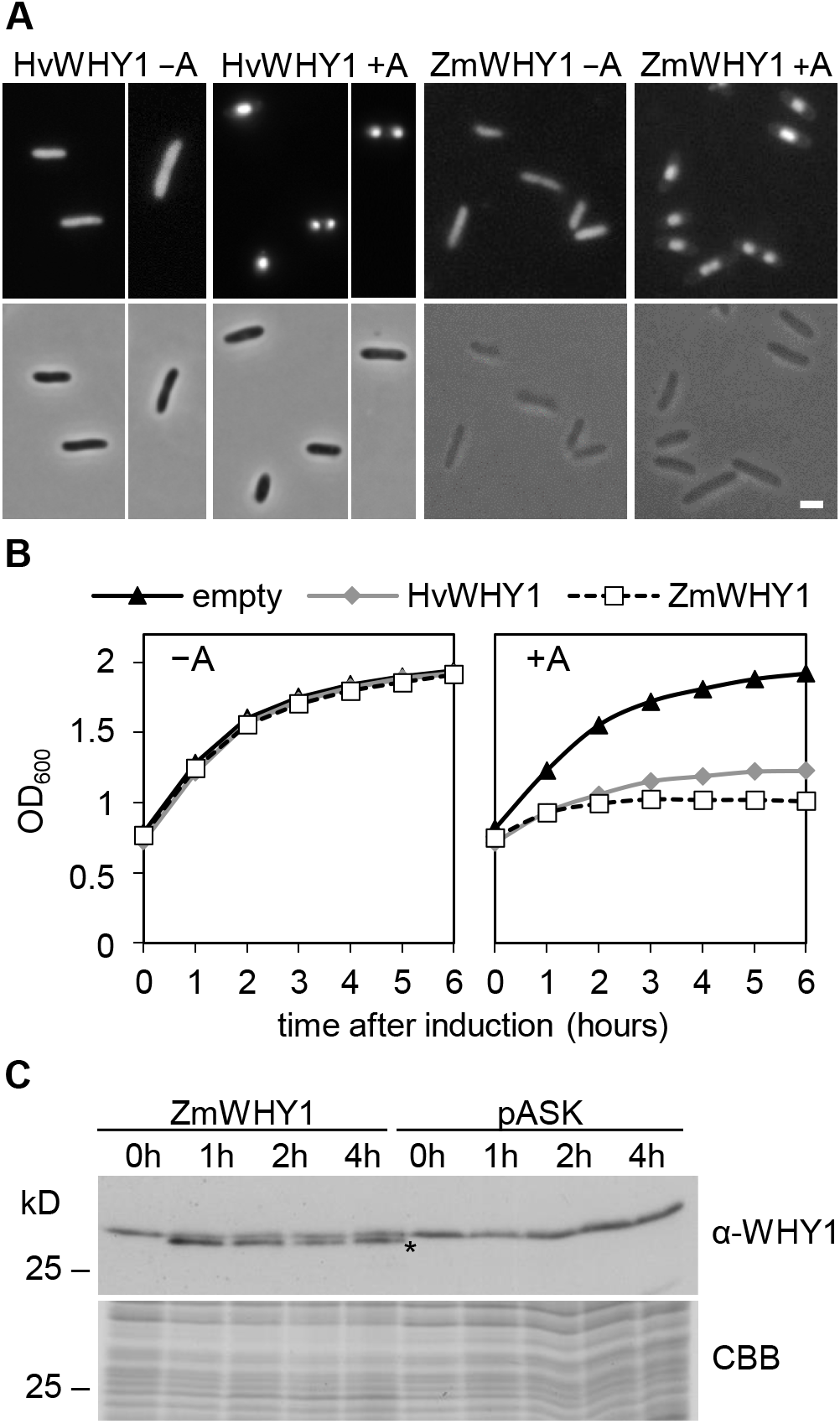
Impact of HvWHIRLY1 and ZmWHIRLY1 on compactness of bacterial nucleoids. *E. coli* DH5α cells were transformed either with the pASK-IBA3 vector containing the gene encoding the mature WHIRLY1 protein of *Hordeum vulgare* (HvWHY1) and *Zea mays* (ZmWHY1), respectively, or the empty pASK-IBA3 vector (empty). −A = no anhydrotetracycline, +A = 200 μg/l anhydrotetracycline. (A) Nucleoids were visualized using DAPI two hours after induction of expression (upper row). Phase contrast images are shown in the lower row. Bar = 2 μm. (B) Growth curves of *E. coli* cells either expressing the empty pASK-IBA3 vector or the *HvWHIRLY1* or *ZmWHIRLY1* coding sequence, respectively. (C) Immunological analysis of bacterial cell lysates expressing the *ZmWHIRLY1* coding sequence zero to four hours after induction of overexpression using an antibody directed against the ZmWHIRLY1 protein. The lower band marked with an asterisk represents the ZmWHIRLY1 protein. A comparable gel was stained with Coomassie Brilliant Blue (CBB).

### The KGKAAL motif including K97 of HvWHIRLY1 does not confer compaction of nucleoids

All WHIRLY proteins share a highly conserved DNA binding motif (DBM) in the WHIRLY domain consisting of the amino acid sequence Lys-Gly-Lys-Ala-Ala-Leu (highlighted with a red bar in Supplementary Figure S2). This motif was shown to confer binding to single stranded DNA (20). The second lysine of the motif (K97 in HvWHIRLY1) (framed in red in Supplementary Figure S2) was shown to be necessary for higher order oligomerization of WHIRLY1 tetramers to 24-mers (21). To investigate whether the DNA binding motif and in particular K97 are required for the nucleoid compacting activity of HvWHIRLY1 in *E. coli*, an *in vitro* mutagenesis was performed. The DNA binding motif of HvWHIRLY1 was either deleted (ΔDBM) or modified by replacement of K97 by an alanine (K97A). Both, cells overexpressing the sequence encoding HvWHIRLY1ΔDBM or HvWHIRLY1K97A contained condensed nucleoids (Figure 2A) and showed a reduced cell growth (Figure 2B) indicating that neither the DNA binding motif nor K97 confer compaction of *E. coli* nucleoids. Immunological analysis of lysates containing the recombinant control HvWHIRLY1 or the mutant proteins, respectively, showed that all proteins are synthesized and soluble in *E. coli* (Figure 2C and Supplementary Figure S3).

**Figure 2.**
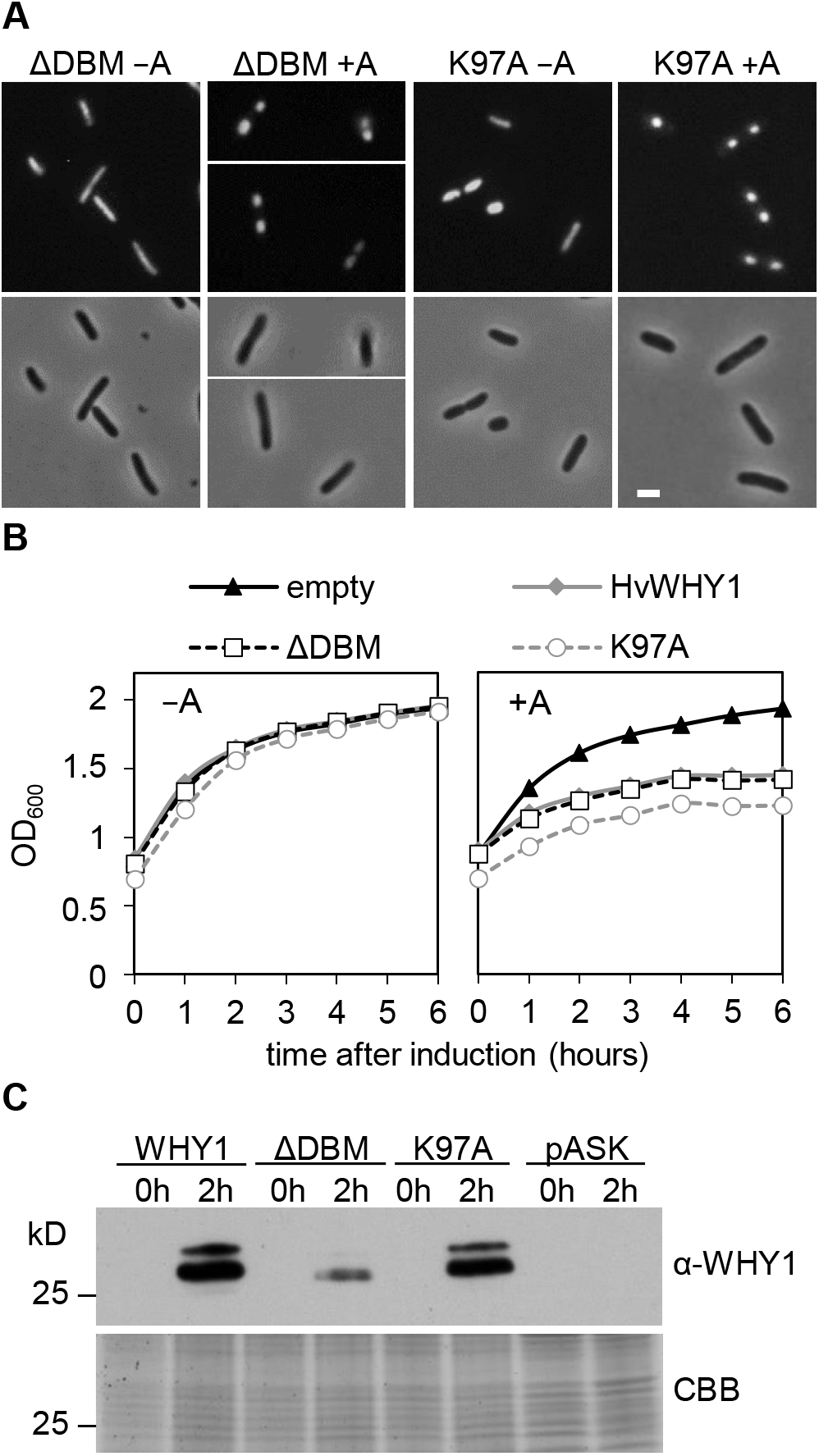
Effect of mutations within the DNA binding motif of HvWHIRLY1 on its compacting activity. *E. coli* DH5α cells were transformed either with the pASK-IBA3 vector containing the gene encoding the mature HvWHIRLY1 protein deleted in the DNA binding motif (ΔDBM), the mature HvWHIRLY1 protein carrying an exchange of lysine 97 against an alanine (K97A) or the empty pASK-IBA3 vector (empty). −A = no anhydrotetracycline, +A = 200 μg/l anhydrotetracycline. (A) Nucleoids were visualized using DAPI two hours after induction (upper row). Phase contrast images are shown in the lower row. Bar = 2 μm. (B) Growth curves of *E. coli* cells either expressing the empty pASK-IBA3 vector, the *HvWHIRLY1* gene or mutated HvWHIRLY1 genes. (C) Immunological analysis of bacterial cell lysates expressing the wild-type or the mutated *HvWHIRLY1* coding sequences shown before and two hours after induction of overexpression, using an antibody directed against the HvWHIRLY1 protein. The bands marked with an asterisk represent the unmodified recombinant protein. A comparable gel was stained with Coomassie Brilliant Blue (CBB).

### Overexpression of *AtWHIRLY1* and *StWHIRLY1* in *E. coli* has no impact on compaction of nucleoids

Arabidopsis chloroplasts contain besides WHIRLY1 the WHIRLY3 protein. To investigate the impact of the two plastidic WHIRLY proteins on compaction of nucleoids, sequences encoding the mature proteins were separately overexpressed in *E. coli*. For comparison, the sequence encoding the mature WHIRLY1 protein of potato, StWHIRLY1, was expressed, too. Neither growth of the bacterial cells (Figure 3A) nor compaction of nucleoids (Supplementary Figure S4) was changed by the overexpression of the sequences, although bacterial cells accumulated the recombinant soluble proteins (Figure 3B and Supplementary Figure S3).

**Figure 3.**
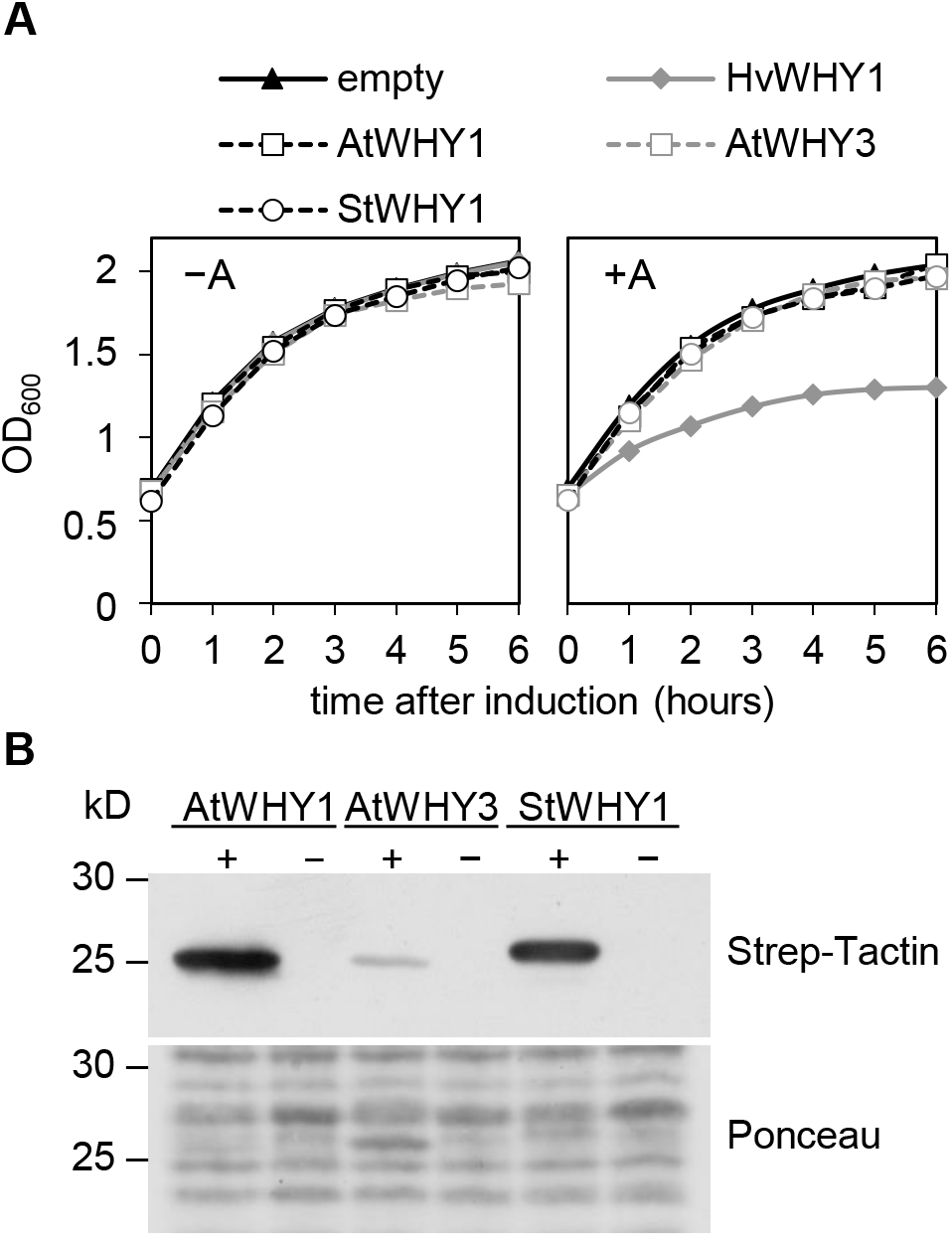
Effect of different WHIRLY1 proteins and the AtWHIRLY3 protein on bacterial cell proliferation and immunological analysis of bacterial cell lysates. (A) Growth curves of *E. coli* cells either expressing the empty pASK-IBA3 vector or plastid *WHIRLY1* and *WHIRLY3* coding sequence from different species. *E. coli* DH5α cells were transformed either with the pASK-IBA3 vector containing the gene encoding the mature WHIRLY1 protein of *Hordeum vulgare* (HvWHY1), the mature WHIRLY1 or WHIRLY3 protein of *Arabidopsis thaliana* (AtWHY1, AtWHY3) and the mature WHIRLY1 protein of *Solanum tuberosum* (StWHY1) or the empty pASK-IBA3 vector (empty). −A = no anhydrotetracycline, +A = 200 μg/l anhydrotetracycline. (B) Immunological analysis of bacterial cell lysates expressing the coding sequences of *AtWHIRLY1* (AtWHY1), *AtWHIRLY3* (AtWHY3) or *StWHIRLY1* (StWHY1) for two hours (+) or of lysates taken from cells without inducing of overexpression collected at the same time (−) using Strep-Tactin®-HRP conjugate. Prior to immunological detection the membrane was stained with Ponceau.

### WHIRLY2 proteins do not affect compaction of nucleoids

The results revealed that nucleoid compaction by HvWHIRLY1 and ZmWHIRLY1 is not due to the WHIRLY domain which is highly conserved in all WHIRLY proteins. Accordingly, mitochondrial WHIRLY2 proteins, neither of barley and maize nor those of Arabidopsis and potato, had any effect on bacterial cell growth and nucleoid organization (Figure 4, Supplementary Figure S3 and Supplementary Figure S5) suggesting that the compacting activity might be restricted to monocot WHIRLY1 proteins.

**Figure 4.**
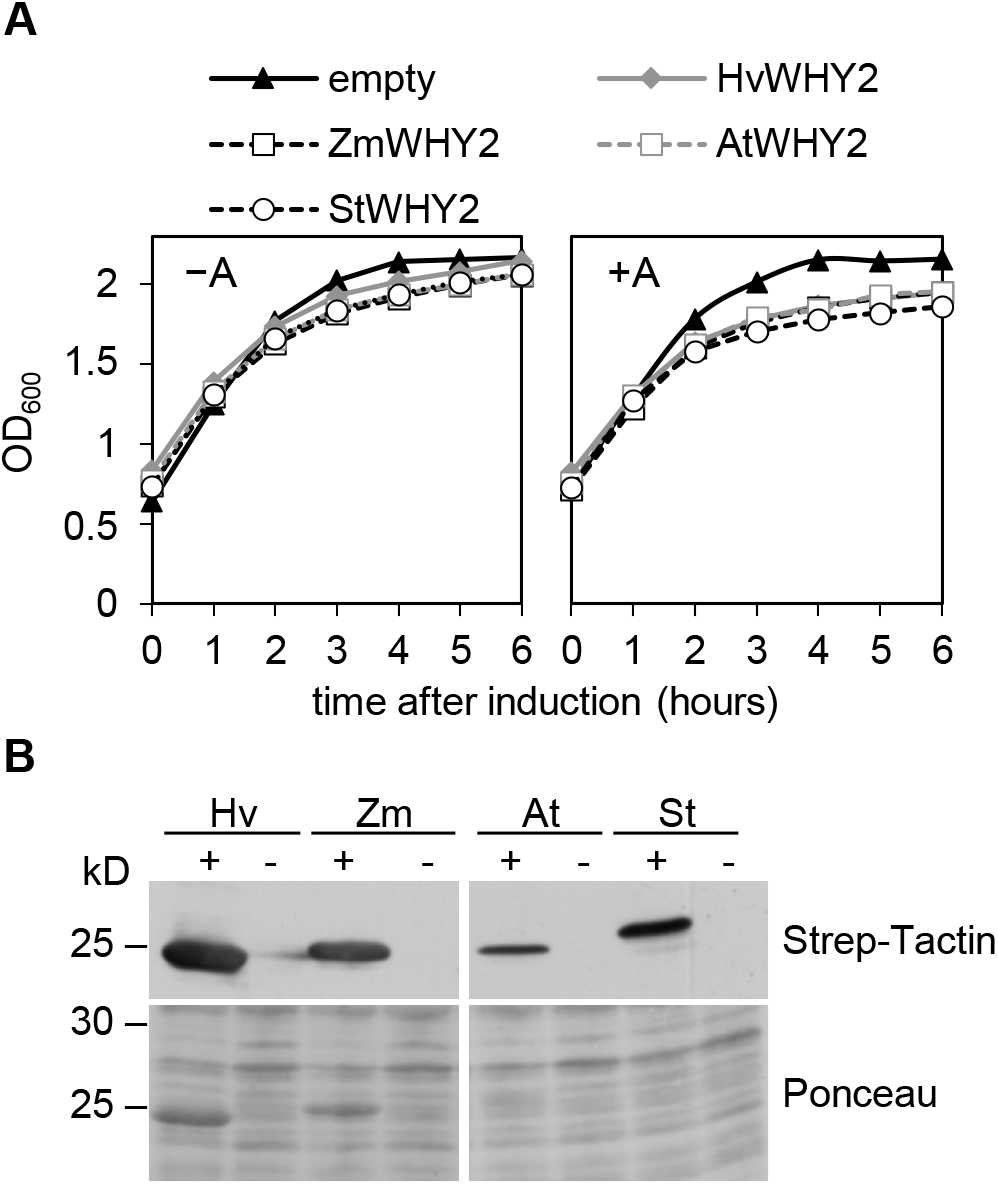
Impact of different WHIRLY2 proteins on bacterial cell proliferation and immunological analysis of bacterial cell lysates. (A) Growth curves of *E. coli* cells either expressing the empty pASK-IBA3 vector or a *WHIRLY2* coding sequence. *E. coli* DH5α cells were transformed either with the empty pASK-IBA3 vector (empty) or the pASK-IBA3 vector containing the gene encoding the mature WHIRLY2 proteins of *Hordeum vulgare* (HvWHY2), *Zea mays* (ZmWHY2), *Arabidopsis thaliana* (AtWHY2) and *Solanum tuberosum* (StWHY2), respectively. −A = no anhydrotetracycline, +A = 200 μg/l anhydrotetracycline. (B) Immunological analysis of lysates from bacterial cells expressing for two hours either the *HvWHIRLY2* (Hv), *ZmWHIRLY2* (Zm), *AtWHIRLY2* (At) or *StWHIRLY1* (St) coding sequences (+) or from cells without induced expression (−) using Strep-Tactin^®^-HRP conjugate. Prior to immunological detection the membrane was stained with Ponceau.

### The PRAPP motif confers compaction of nucleoids

A sequence alignment with selected WHIRLY1 proteins of the Poaceae family showed a conserved region located between the plastid target peptide and the WHIRLY domain (Figure 5A) that is neither present in WHIRLY1 proteins of dicots nor in WHIRLY2 proteins of both dicots and Poaceae (Supplementary Figure S6). To investigate whether this region of the sequence is required for compaction of bacterial nucleoids, a deletion of residues 42-60 (Δ42-60) of the mature HvWHIRLY1 was performed by *in vitro* mutagenesis (Figure 5A, framed sequence). Neither growth of bacterial cells (Figure 5B) nor nucleoid compaction (Supplementary Figure S7A) was affected by expression of HvWHIRLY1 lacking residues 42-60. This result revealed that the amino acid residues responsible for compaction of nucleoids are contained in this region of the sequence. The accumulation of the recombinant HvWHIRLY1Δ42-60 was verified with immunological analysis (Supplementary Figure S7B).

**Figure 5.**
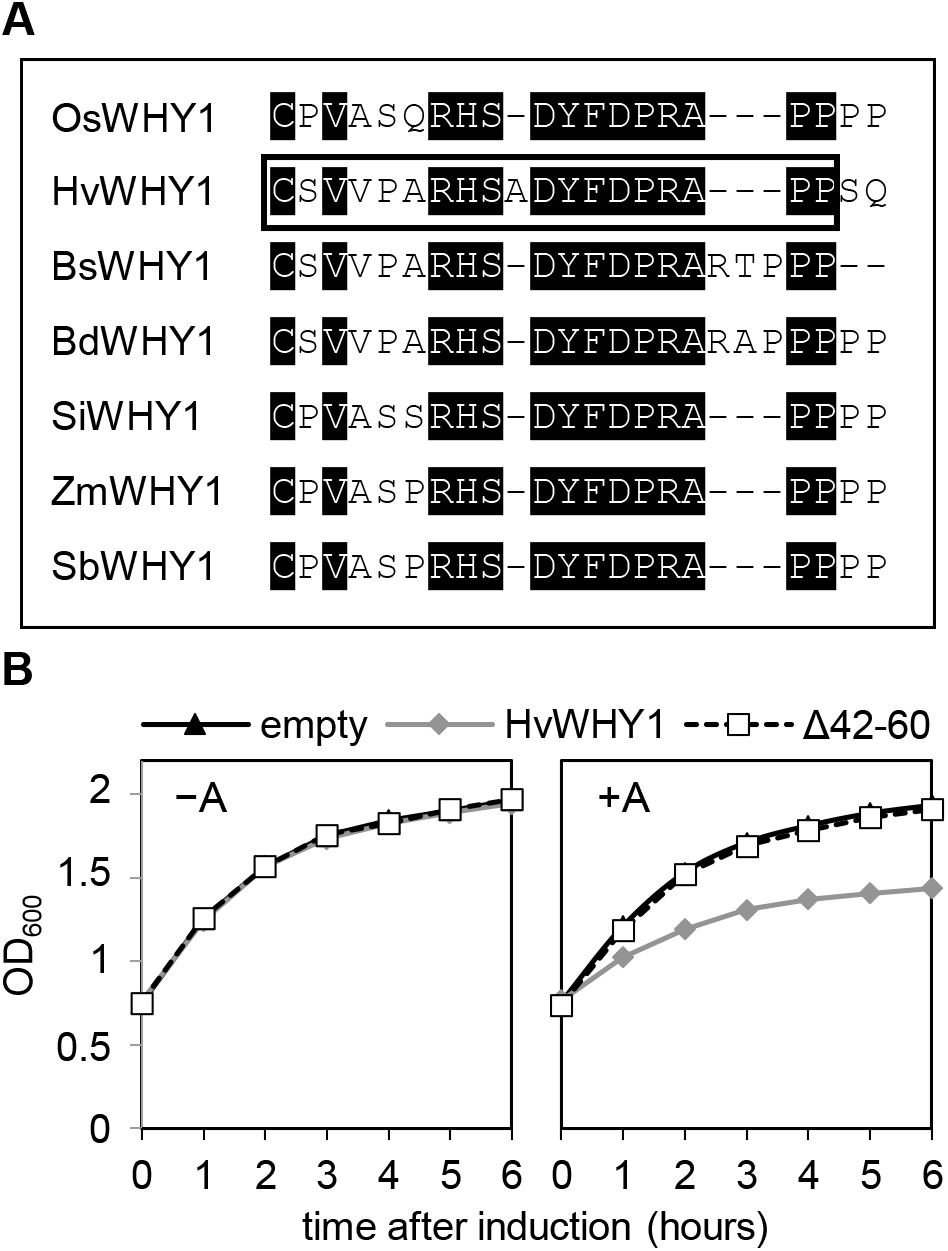
Impact of mutated *HvWHIRLY1Δ42-60* sequence on bacterial cell proliferation. (A) Alignment of the conserved N-terminal part of WHIRLY1 proteins from monocots. The framed residues 42-60 were deleted by *in vitro* mutagenesis. (B) Growth curves of *E. coli* cells either expressing the empty pASK-IBA3 vector (empty), the *HvWHIRLY1* (HvWHY1) coding sequence or the mutated *HvWHIRLY1Δ42-60* (Δ42-60) coding sequence. −A = no anhydrotetracycline, +A = 200 μg/l anhydrotetracycline.

The sequence covered by residues 42-60 of HvWHIRLY1 contains two arginines, which are positively charged and hence might interact with nucleic acids. They are combined with several prolines which are known to form stiff ring structures leading to kinks in the tertiary structure of proteins (52). The amino acid motif Pro-Arg-Ala-Pro-Pro (PRAPP) which is found within residues 42-60 is shared by a subset of WHIRLY1 proteins (23) including also maize and barley and is likely responsible for the compaction of nucleoids in these plants.

To investigate the role of the prolines and the arginine for the compacting activity of WHIRLY1, three mutations were introduced in barley *WHIRLY1* using *in vitro* mutagenesis. The residues 56-60 (ΔPRAPP) were deleted, the three prolines P56, P59 and P60 (P/A) were altogether replaced by alanines and the single arginine R57 was replaced by an alanine (HvR57A) (Figure 6). Although the recombinant mutant proteins were shown to be soluble (Supplementary Figure S3) and accumulated in higher amounts than the control HvWHIRLY1 protein two hours after induction of overexpression (Figure 7B), neither had an effect on cell proliferation (Figure 7A) nor on the morphology of nucleoids of the cells was observed (Supplementary Figure S8). When the arginine R63 of the ZmWHIRLY1 protein corresponding to R57 in the PRAPP motif of HvWHIRLY1, was replaced by an alanine (ZmR63A) (Figure 6) the compaction activity was lost, too (Figure 7 and Supplementary Figure S8). These results indicate that the PRAPP motif is responsible for the compacting activities of HvWHIRLY1 and ZmWHIRLY1.

**Figure 6.**
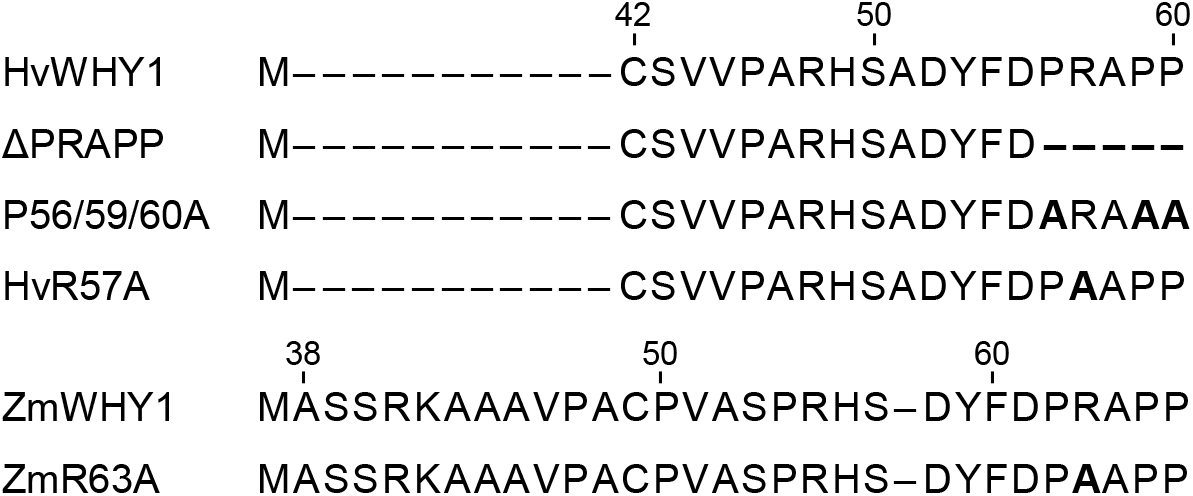
Partial amino acid sequences of mutated HvWHIRLY1 and ZmWHIRLY1 examined for nucleoid compaction activity. The mature HvWHIRLY1 amino acid sequence used for heterologous expression experiments starts at residue 42 behind an artificial methionine and the mature ZmWHIRLY1 amino acid sequence starts at residue 38. Numbers indicate the positions of the amino acids in the mature WHIRLY1 proteins. Bold symbols show amino acid changes or deletions introduced by *in vitro* mutagenesis into the WHIRLY1 protein.

**Figure 7.**
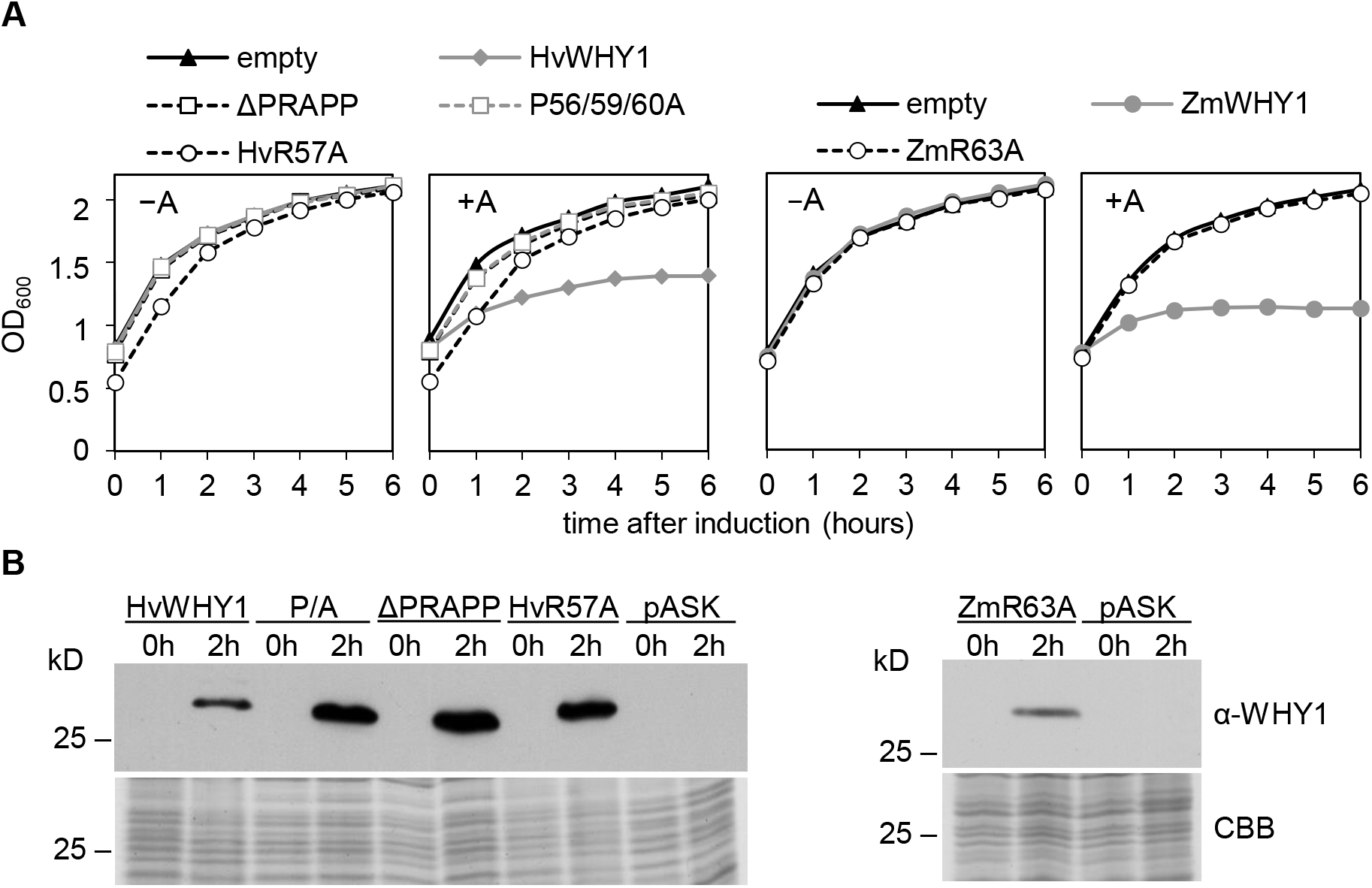
Effect of altered PRAPP motifs on bacterial cell proliferation and immunological analysis of bacterial cell lysates. (A) Growth curves of *E. coli* cells either expressing the empty pASK-IBA3 vector (empty), the *HvWHIRLY1* (HvWHY1), the *ZmWHIRLY1* (ZmWHY1), the mutated *HvWHIRLY1*ΔPRAPP (ΔPRAPP), *HvWHIRLY1P56/59/60A* (P/A), *HvWHIRLY1R57A* (R57A) or ZmWHIRLY1R63A (R63A) coding sequence. −A = no anhydrotetracycline, +A = 200 μg/l anhydrotetracycline. (B) Immunological analysis of bacterial cell lysates expressing the *HvWHIRLY1* (HvWHY1), *HvWHIRLY1*ΔPRAPP (ΔPRAPP), *HvWHIRLY1P56/59/60A* (P56/59/60A), *HvWHIRLY1R57A* (HvR57A) or *ZmWHIRLY1R63A* (ZmR63A) coding sequence before and two hours after induction of overexpression using an antibody directed against the HvWHIRLY1 protein and ZmWHIRLY1 protein, respectively. A comparable gel was stained with Coomassie Brilliant Blue (CBB).

### HvWHIRLY1 dependent nucleoid compaction effects transcript levels of genes in *E. coli* after overexpression of HvWHIRLY1

To investigate the effect of HvWHIRLY1 dependent nucleoid compaction on transcript levels of *E. coli* genes, a qRT-PCR analysis was performed. The relative transcript levels of five genes encoding proteins involved in DNA replication (*dnaA*) and RNA transcription (*rpoA*), cell division (*ftsZ*) and nucleoid organization (*hupA* and *hns*) were analysed in HvWHIRLY1 and ΔPRAPP producing cells in comparison to cells containing the empty pASK-IBA3 vector (Figure 9). *idnT* and *cysG*, known as stable *E. coli* genes to study transcriptional alterations due to recombinant protein accumulation (Zhou et al. 2011), were used as reference genes. The qRT-PCR data show that the WHIRLY1 dependent nucleoid compaction resulted in an enhanced accumulation of *dnaA* and *rpoA* transcripts, whereas *ftsZ*, *hupA* and *hns* transcripts were reduced. Cells producing HvWHIRLY1ΔPRAPP did not show such alterations, the transcript levels were rather similar to cells containing the empty vector or to cells not expressing any *WHIRLY1* coding sequence. This result indicated that the altered transcript levels are caused by the nucleoid compaction by WHIRLY1 and not just by the accumulation of recombinant proteins. The reference gene *idnT* also supports this result because its transcript level was comparable in all tested cells and thus is not affected by recombinant protein accumulation.

### Addition of the PRAPP motif enables AtWHIRLY1 to condense *E. coli* nucleoids

The mutations introduced in *HvWHIRLY1* and *ZmWHIRLY1*, respectively, resulted in the elimination of the compacting activity of these two proteins indicated by a normal cell growth curve and loosely packed nucleoids. To examine whether the addition of the PRAPP motif in the amino acid sequence of AtWHIRLY1 can confer compacting activity to AtWHIRLY1, a chimeric protein consisting of the mature AtWHIRLY1 protein and the PRAPP motif was generated by *in vitro* mutagenesis. An appropriate insertion site for PRAPP was identified with a sequence alignment of the HvWHIRLY1 and AtWHIRLY1 protein (Figure 8A). The accumulation of the chimeric AtWHIRLY1+PRAPP protein resulted in an even greater reduction of the *E. coli* cell proliferation rate than accumulation of HvWHIRLY1 (Figure 8B). Furthermore, the cells contained densely packed nucleoids (Supplementary Figure S9). Similar to HvWHIRLY1, recombinant AtWHIRLY1+PRAPP accumulated in smaller quantities in the cells than the recombinant AtWHIRLY1 protein (Figure 8C) suggesting that WHIRLY proteins that have an effect on bacterial nucleoid condensation are expressed at lower levels due to an altered accessibility of the bacterial genome compared to proteins that do not possess a compacting activity.

**Figure 8.**
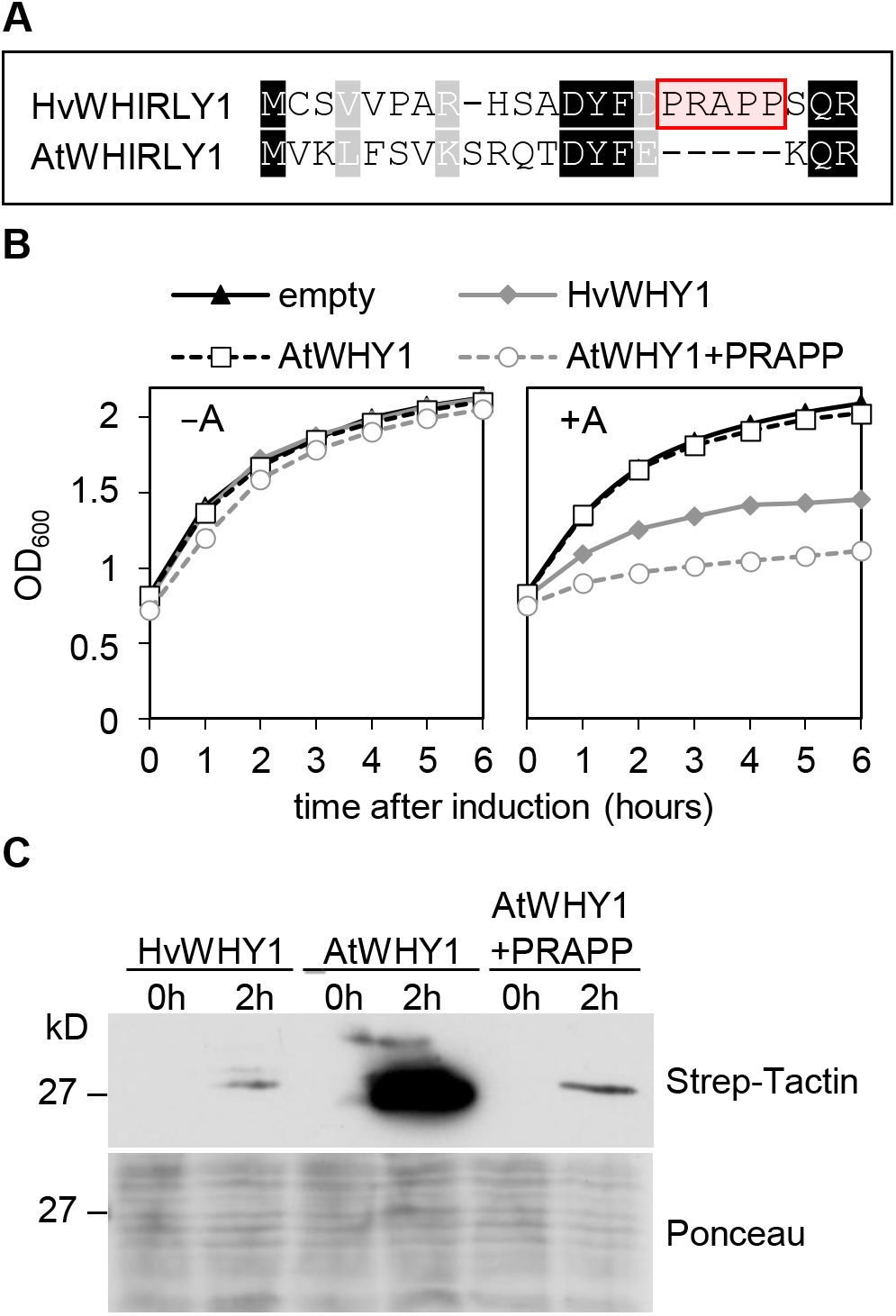
Effect of a chimeric AtWHIRLY1+PRAPP protein on *E. coli*. (A) Alignment of the N-terminal region of the WHIRLY1 protein from *Hordeum vulgare* and *Arabidopsis thaliana*. The framed PRAPP motif was introduced in AtWHIRLY1 protein at the indicated position. (B) Growth curves of *E. coli* cells either expressing the empty pASK-IBA3 vector (empty), *HvWHIRLY1* (HvWHY1) or *AtWHIRLY1* (AtWHY1) coding sequence or the *AtWHIRLY1+PRAPP* (AtWHY1+PRAPP) coding sequence. −A = no anhydrotetracycline, +A = 200 μg/l anhydrotetracycline. (C) Immunological analysis of bacterial cell lysates expressing the *HvWHIRLY1* (HvWHY1), *AtWHIRLY1* (AtWHY1) or *AtWHIRLY1+PRAPP* (AtWHY1+PRAPP) coding sequence before and two hours after induction of overexpression using Strep-Tactin®-HRP conjugate. Prior to immunological detection the membrane was stained with Ponceau.

### Overexpression of chimeric *AtWHIRLY1+PRAPP* has no effect on the nucleoid structure in chloroplasts of Arabidopsis

To investigate the compacting activity of AtWHIRLY1+PRAPP *in planta*, *A. thaliana* ecotype Col-0 was stably transformed with the binary vector pB2GW7,0 encoding the coding sequence of *AtWHIRLY1* carrying the PRAPP motif and an HA tag at the C-terminus (Figure 10A). Although all tested AtWHIRLY1+PRAPP:HA overexpression lines accumulated the HA-tagged AtWHIRLY1+PRAPP protein in the plastids (Figure 10B), no altered phenotype was observed in comparison to an AtWHIRLY1:HA overexpression line (OE) (53) under standard growth conditions (Figure 10C).

**Figure 9.**
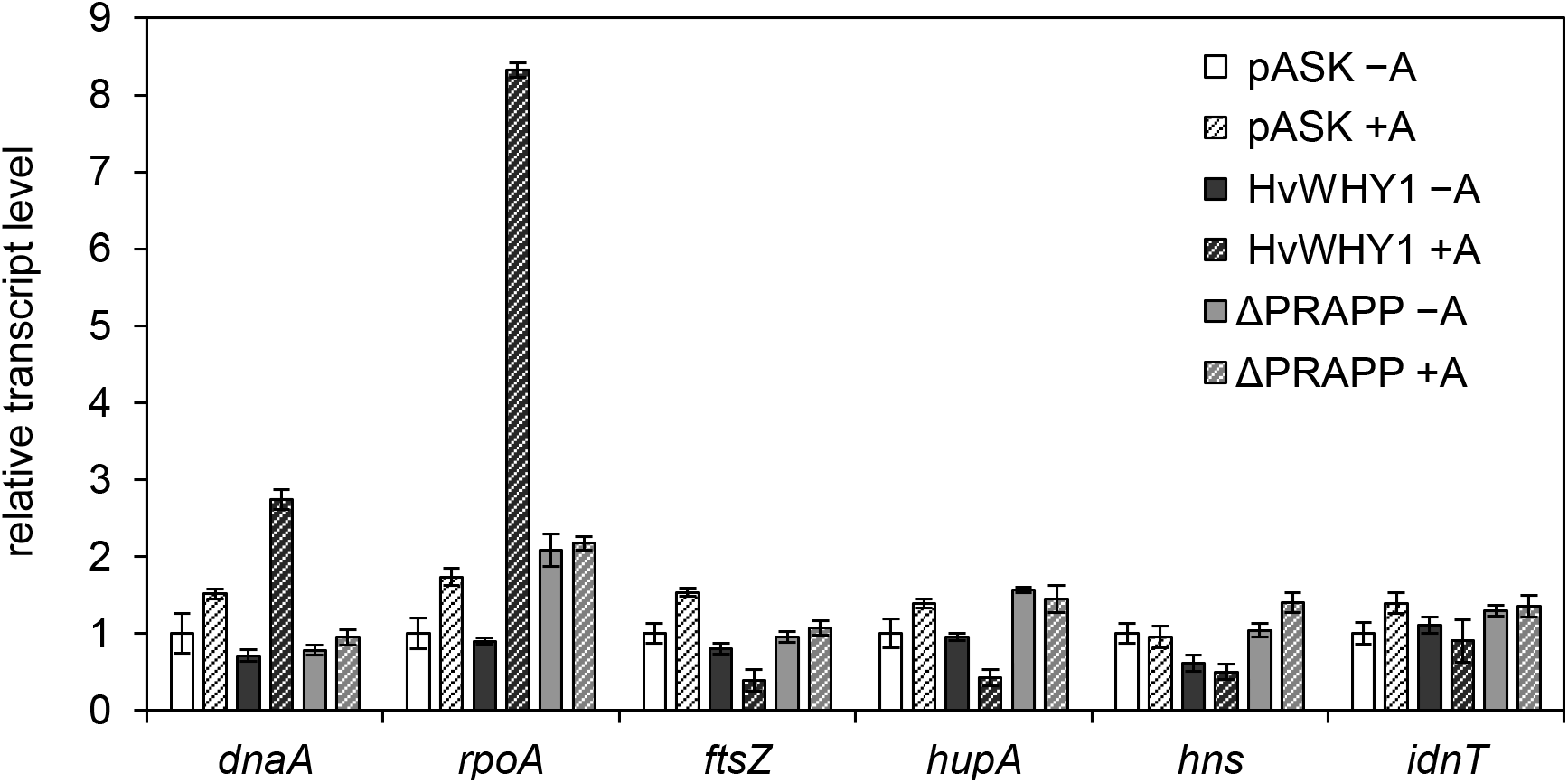
Transcript levels of genes in *E. coli* cells two hours after induction of overexpression of *HvWHIRLY1* and *HvWHIRLY1ΔPRAPP*. RNA was isolated two hours after induction from bacterial cells containing the empty pASK-IBA3 vector (pASK) or expressing either the *HvWHIRLY1* (*HvWHY1*) or *HvWHIRLY1ΔPRAPP* (*ΔPRAPP*) coding sequences (+A) or from cells without induced expression (−A). Relative transcription levels were normalized to the *cysG* transcript and data are shown in relation to *E. coli* cells containing the empty vector pASK-IBA3 without induction. *dnaA* = catalytic subunit of DNA-dependent DNA polymerase III, *rpoA* = DNA-dependent RNA polymerase, *ftsZ* = cell division protein that forms contractile Z ring, *hupA* = histone-like DNA-binding protein alpha, *hns* = DNA-binding protein HNS, *idnT* = Gnt-II system L-idonate transporter.

**Figure 10.**
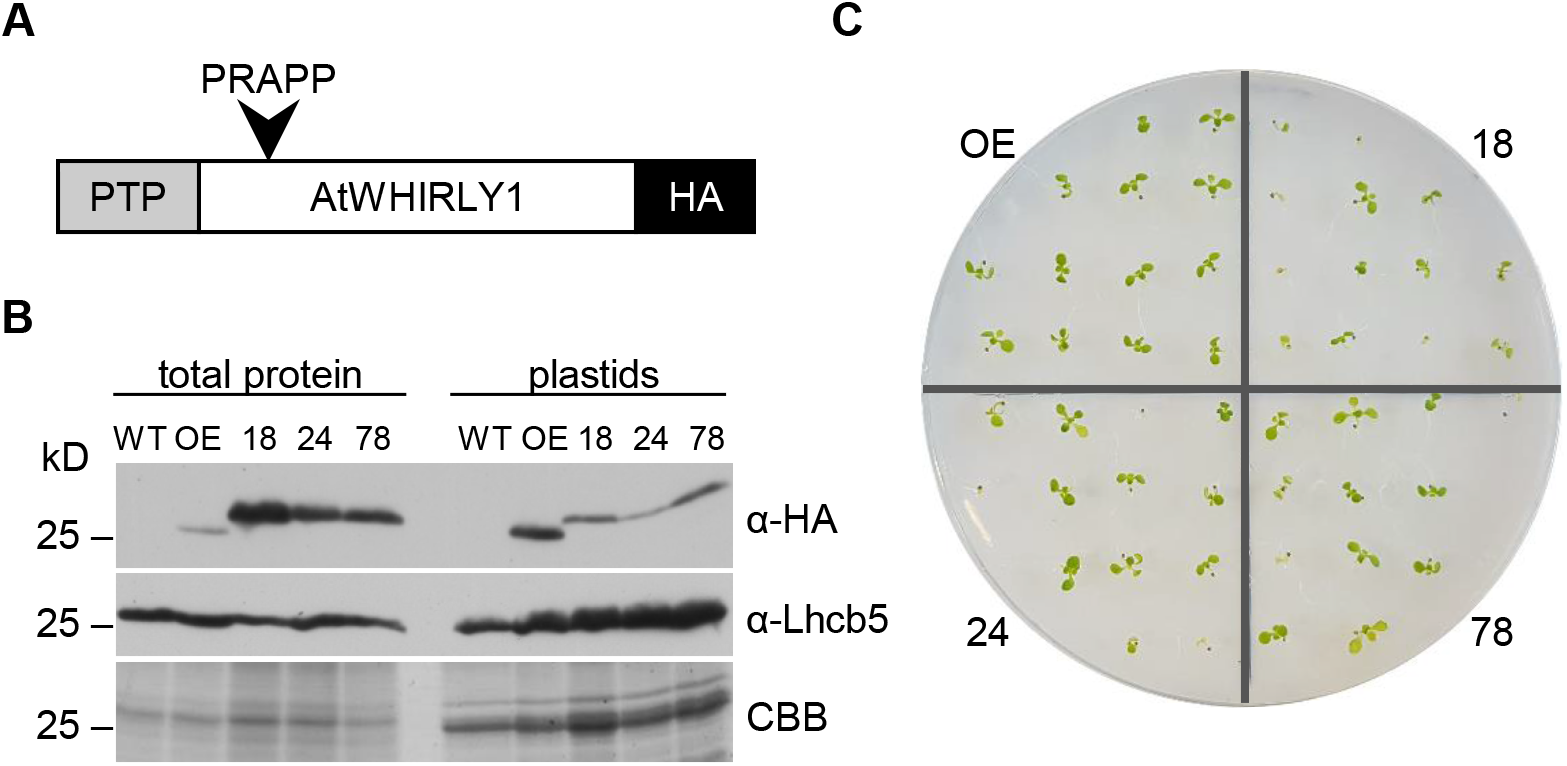
Characterisation of AtWHIRLY1+PRAPP:HA overexpressing lines. (A) Scheme of the construct used for stable transformation of the HA-tagged AtWHIRLY1+PRAPP. PTP = plastid target peptide, HA = hemagglutinin tag. (B) Immunological analysis of the overexpressing lines. 15 μg total protein extracts and 15 μg plastids (α-HA and CBB) or 5 μg plastids (α-Lhcb5) were subjected to SDS-PAGE. Detection of HA-tagged AtWHIRLY1+PRAPP was done using an HA tag antibody. As a control for the plastid fraction the Lhcb5 antibody was used. A comparable gel was stained with Coomassie Brilliant Blue (CBB). (C) Phenotype analysis of the AtWHIRLY1+PRAPP:HA overexpressing lines 18, 24, 78 and an AtWHIRLY1:HA overexpressing line (OE) (described in Isemer et al. 2012, named pnWHIRLY1:HA) 12 days after sowing. Surface sterilized seeds were sown on MS agar plates supplemented with Basta^®^ to a final concentration of 1:50,000 for selection.

To analyse whether accumulation of AtWHIRLY1+PRAPP:HA leads to an altered nucleoid morphology, cross-sections from the first true leaves of plants 14 days after sowing were fixed by formaldehyde and were stained with SYBR Green I. This analysis revealed that the added PRAPP motif did not have any obvious effect on the nucleoid structure in Arabidopsis because no difference could be observed in AtWHIRLY1+PRAPP:HA plants in comparison to AtWHIRLY1:HA plants (Supplementary Figure S10). However, the fluorescence signals from nucleoids in all overexpression lines were more distinct than in the WT indicating that the overexpression of either *AtWHIRLY1:HA* and *AtWHIRLY1+PRAPP:HA* led to an altered morphology of plastid nucleoids independent of the PRAPP motif.

## DISCUSSION

The major part of the WHIRLY proteins is highly conserved and has been called WHIRLY domain, consisting of eight ß-sheets and three alpha helices (23). The N- and C-terminal parts are more variable. N-terminal parts contain in some of the WHIRLY proteins potential transactivation domains (23). Such diverse transactivation sequences in different species, rich in serines, prolines or glutamines, indicate functional divergence (54).

Due to lacking sequence information of the N-terminal part at the time, the HvWHIRLY1 together with WHIRLY2 proteins of maize, wheat and rice were assigned to those WHIRLY proteins lacking a transactivation domain (MSII) while the WHIRLY1 proteins of rice, maize, sorghum and wheat were grouped together in the monocot subfamily of proteins having a polyproline activation domain (MSI) (23). The complete sequence of HvWHIRLY1 available after sequencing of the complete barley genome (55) does possess a proline-rich N-terminus like the other monocot WHIRLY1 sequences in which the PRAPP motif responsible for the compacting activity of WHIRLY1 proteins is located. The diverse motifs besides the highly homologous WHIRLY domain enabled plants in different groups to use these proteins for diverse functions.

### Evolution of the PRAPP motif in WHIRLY1 proteins

WHIRLY1 proteins with PRAPP motifs are found in many members of the Poaceae family. Among them are wheat, rice and sorghum. Besides the PRAPP motif, also a PRAPP-like motif which is interrupted by two additional amino acids is found in two WHIRLY1 proteins in the Poaceae i.e., of *Brachypodium distachyon* (*B. distachyon*, BdWHIRLY1) and *Brachypodium sylvaticum* (*B. sylvaticum*, BsWHIRLY1) thus is similar to the WHIRLY1 sequence of *Ananas comosus* (*A. comosus*, AcWHIRLY1) which is interrupted by four additional amino acids, too (Figure 11). *A. comosus* is a member of the Bromeliaceae which are basal Poales (56) and diverged earlier than the Poaceae (56,57). Hence, the PRAPP-like motif of *A. comosus*, rich in prolines, may represent an early precursor of the PRAPP motif of the Poaceae WHIRLY1 proteins (Figure 11). However, whether the PRAPP-like motif can confer compacting activity to the WHIRLY1 proteins of *A. comosus*, *B. distachyon* and *B. sylvaticum* remains to be shown. Although WHIRLY1 proteins found in species belonging to the order of Arecales and Zingiberales do not possess a PRAPP motif, they have proline and arginine residues at the N-terminal part of the protein sequence corresponding to the region where PRAPP is found in the WHIRLY1 proteins of the Poaceae (Figure 11). In contrast, no proline or arginine residues are present in the WHIRLY1 protein of *Zostera marina* (*Z. marina*, ZmaWHIRLY1) belonging to the Alismatales, the oldest order of monocots here mentioned (58). This finding indicates that proline and arginine residues evolved in WHIRLY1 proteins after the divergence of Alismatales in the Arecales, Zingiberales and Poales and that the formation of the PRAPP motif is restricted exclusively to the Poaceae family.

**Figure 11.**
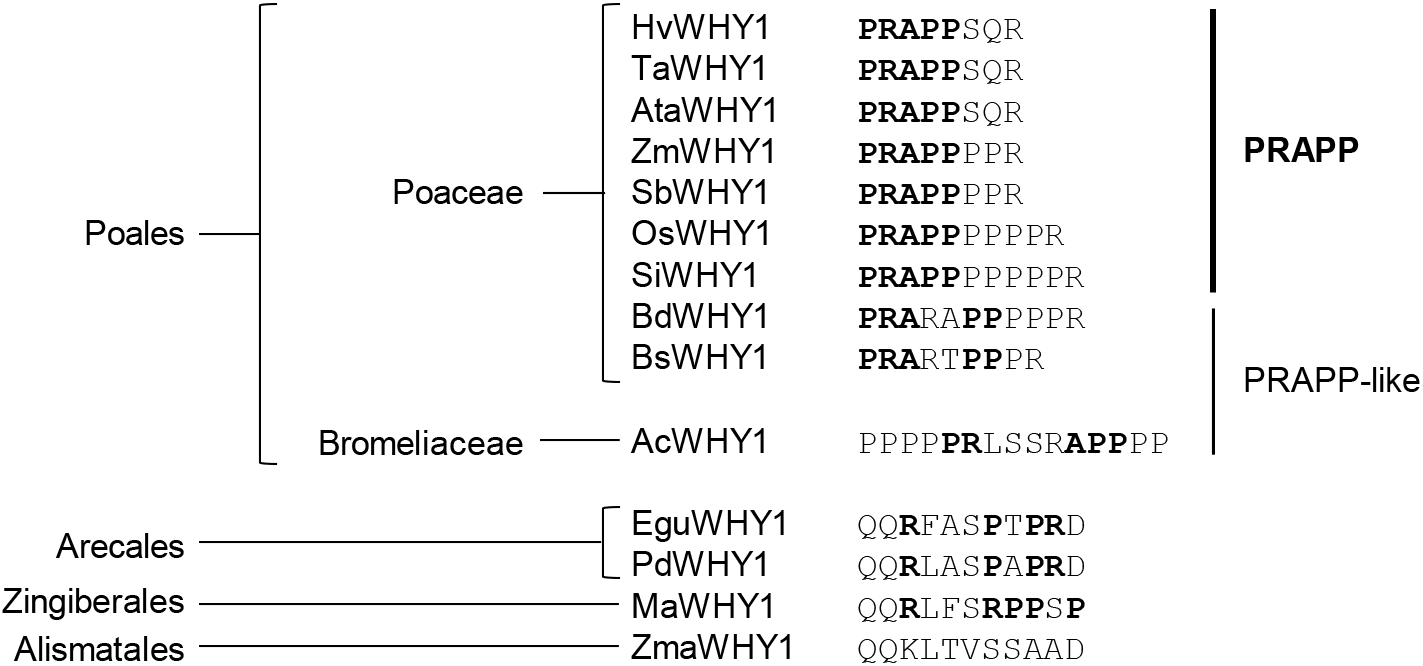
Sequences with PRAPP and PRAPP-like motifs in monocot WHIRLY1 proteins. The PRAPP motif, single proline and arginine residues are shown in bold. The following protein sequences were used (name, organism, GenBank accession number): AcWHY1, *Ananas comosus*, OAY85479.1; AtaWHY1, *Aegilops tauschii*, XM_020331194.1; BdWHY1, *Brachypodium distachyon*, XP_003557198; BsWHY1, *Brachypodium sylvaticum*, ABL85062; EguWHY1, *Elaeis guineensis*, XP_010915142.1; HvWHY1, *Hordeum vulgare*, BAJ96655; MaWHY1, *Musa acuminata*, XP_009380437.1; OsWHY1, *Oryza sativa*, BAD68418; PdWHY1, *Phonenix dactylifera*, XP_008809721.1; SbWHY1, *Sorghum bicolor*, XP_002436467; SiWHY1, *Setaria italica*, XP_004964537; TaWHY1, *Triticum aestivum*, AK454665.1; ZmWHY1, *Zea mays*, NP_001123589; ZmaWHY1, *Zostera marina*, KMZ68486.1.

### Proline and arginine residues facilitate the interaction of bacterial NAPs with dsDNA

The PRAPP motif was shown to be responsible for the compacting activity of WHIRLY1 proteins in barley and maize. However, overexpression of *ZmWHIRLY1* in *E. coli* caused a stronger inhibition of bacterial cell growth than the expression of *HvWHIRLY1*. The difference could be due to two additional proline residues (PRAPP**PP**). Prolines are known to form stiff ring structures leading to kinks within the tertiary structure of proteins (52). The additional prolines of the PRAPP**PP**motif could further strengthen the interaction of WHIRLY1 with DNA thus resulting in a stronger condensation of bacterial nucleoids. Mutation of the highly conserved DNA binding motif (ΔDBM) within the WHIRLY1 domain had no impact on condensation of nucleoids in *E. coli* indicating that the ability to bind to single stranded DNA is not necessary for the condensation of nucleoids by WHIRLY1 proteins. Electrophoretic mobility shift assays furthermore showed that a recombinant ZmWHIRLY1 protein is able to bind double stranded DNA and to bind plastid DNA in a sequence non-specific manner (25). Hence, it is supposed that the PRAPP-containing WHIRLY1 proteins can interact with double stranded DNA without any sequence specificity like it is the case for the histones in eukaryotic nuclei (59) and the histone-like proteins of bacteria. Prolines are also known to be involved in the structural organization of DNA by HU and IHF, two bacterial NAPs that induce bending of double stranded DNA in *E. coli*, each of them acting as a (hetero-)dimer (60). Both HU and IHF interact via two β-ribbon arms (one β-ribbon arm from each subunit) with the minor groove of the DNA. At the tip of each β-ribbon arm a proline intercalates between base pairs inducing two kinks in the DNA resulting in bending (60,61). Moreover, it has been shown that two arginine residues located two and five amino acids N-terminally of the proline, respectively, interact via hydrogen bonds with base pairs of the minor groove of the DNA (61). It remains to be shown whether the prolines and arginine of the WHIRLY1 PRAPP motif act on DNA in a similar manner as those of IHF and HU. Compaction of bacterial nucleoids depends on supercoiling, bending and wrapping of DNA (62). All DNA associated activities are performed by dimers and higher oligomeric structures of the corresponding NAPs. The importance of dimerization and oligomerization of WHIRLY1 for chloroplast nucleoid compaction remains to be studied *in planta* with mutated forms of the protein.

### The relationship between WHIRLY1 and other eukaryotic architectural proteins of plastid nucleoids of higher plants

In contrast to WHIRLY1 of barley and maize the WHIRLY1 proteins of Arabidopsis and potato did not show a nucleoid compacting activity in *E. coli*. Nevertheless, plastid nucleoids are well compact in chloroplasts of Arabidopsis despite the lack of WHIRLY1. In a previous study it has been shown that Arabidopsis has a SWIB-4 protein with a histon H1 like motif able to compact nucleoids in *E. coli* cells (2). It is likely that SWIB-4 is responsible for compaction of nucleoids in chloroplasts. So far, no mutant lacking AtSWIB-4 was available for an analysis of nucleoid morphology.

The first nucleoid associated protein shown to induce compaction of plastid DNA is the 68 kDa sulfite reductase (SIR) whose primary function in sulfur assimilation is not related to DNA (63). SIR like WHIRLY1 binds to ssDNA and dsDNA without any sequence specificity (25,64). The compacting activity of SIR is mediated by a C-terminally encoded peptide (CEP) (65). This CEP is predicted to form a bacterial ribbon-helix-helix DNA binding motif that could facilitate interaction with plastid nucleoids (65). SIR of *Nicotiana tabacum* (NbSIR), *Pisum sativum* (PsSIR) and *Glycine max* (GmSIR) can act as nucleoid compacting proteins. Recombinant NbSIR was shown to condense nucleoids in *E. coli* (66), whereas PsSIR and GmSIR were shown to condense chloroplast DNA *in vitro* (63,64). In contrast, the dsDNA binding ability of maize SIR lacking the CEP was shown to be weaker than that of the pea SIR (64). Furthermore maize and pea SIR also differ in their ability to compact chloroplast DNA *in vitro* indicating that SIR proteins need the CEP for nucleoid compaction. Indeed, several members of the Brassicaceae and the Poaceae such as maize and Arabidopsis possess SIR without a CEP (65). Although CEP was found to be necessary for nucleoid localization and compaction, the addition of CEP at the C-terminus to the Arabidopsis SIR (AtSIR) merely resulted in a co-localization of AtSIR+CEP with plastid nucleoids but did not affect the compaction of plastid DNA (65).

### AtWHIRLY1 with an inserted PRAPP motif has no impact on the compactness of plastid nucleoids

Overexpression of a construct encoding AtWHIRLY1 with a PRAPP motif inserted induced compaction of nucleoids in *E. coli*. In contrast, no enhanced compaction of plastid nucleoids was observed in transgenic Arabidopsis lines expressing the chimeric *AtWHIRLY1+PRAPP:HA* compared to an *AtWHIRLY1:HA* overexpression line. The result is in accordance with the green phenotype of the *why1* mutant (Salk_023713) (30) and also the *why1 why3* double mutant (35) which under standard growth conditions don’t have an altered nucleoid morphology in comparison to the wild type (data not shown). In contrast to the *why* mutants of Arabidopsis, in the maize *why1* mutants (25) and the RNAi-W1 lines of barley (41) chloroplast development is severely affected. This is a consequence of impaired ribosome formation which is known to depend on nucleoids serving as a platform (40).

The lacking impact of PRAPP on the nucleoid compaction in Arabidopsis is also interesting with regard to the findings of (65) showing that the addition of CEP to the Arabidopsis SIR protein had also no visible effect on the structural organization of plastid nucleoids. Obviously, both motifs don’t function in all plants indicating that different plants have different proteins for compaction of plastid nucleoids. Neither WHIRLY1 nor SIR have a function in compacting plastid nucleoids in Arabidopsis. This function might be resumed by SWIB-4 whose overexpression in *E. coli* had a compacting effect on the bacterial nucleoids (2). Taken together the data indicate an intriguing divergence of nucleoid architectural proteins in angiosperms. As shown by the unpacked nucleoid morphology of chloroplasts in transgenic barley plants deficient in WHIRLY1 (27) and the results of this study, WHIRLY1 seemingly is the principle architectural protein of plastid nucleoids within the family of Poaceae.

WHIRLY proteins are multifunctional proteins due to different putative transactivation regions at the N-terminus (23). Obviously, WHIRLY1 proteins characterized by a PRAPP have acquired an additional function as plastid nucleoid architectural protein within the family of Poaceae. This function is not dependent on the highly conserved WHIRLY domain shared by all WHIRLY proteins.

This result revealed that the loss of WHIRLY1 had no influence on the nucleoid organization in Arabidopsis being in accordance with the heterologous expression analyses in *E. coli*, thus indicating that the compacting activity is restricted to WHIRLY1 proteins with PRAPP motif exclusively found in the Poaceae.

## Supporting information

Supplement complete

## SUPPLEMENTARY DATA

Supplementary Data are available at NAR online.

## ACKNOWLEDGEMENT

We thank Dr. Götz Hensel (IPK Gatersleben) for discussion and critical reading of the manuscript. Anke Schäfer (Institute of Botany, CAU Kiel) is thanked for excellent technical assistance. We thank the Institute of Clinical Molecular Biology in Kiel for providing Sanger sequencing as supported in part by the DFG Cluster of Excellence “Inflammation at Interfaces” and “Future Ocean”. We thank the technicians T. Naujoks and C. Noack for technical support. The Central Microscopy of the Biology Centre (CAU Kiel) is thanked for providing microscopic facilities.

## FUNDING

This work was supported by German Research Foundation (DFG) [KR1350/19-1].

## Notes

### Competing Interest Statement

The authors have declared no competing interest.

