## Supplement complete for "WHIRLY1 of barley and maize share a PRAPP motif conferring nucleoid compaction"

**Supplementary Figures**

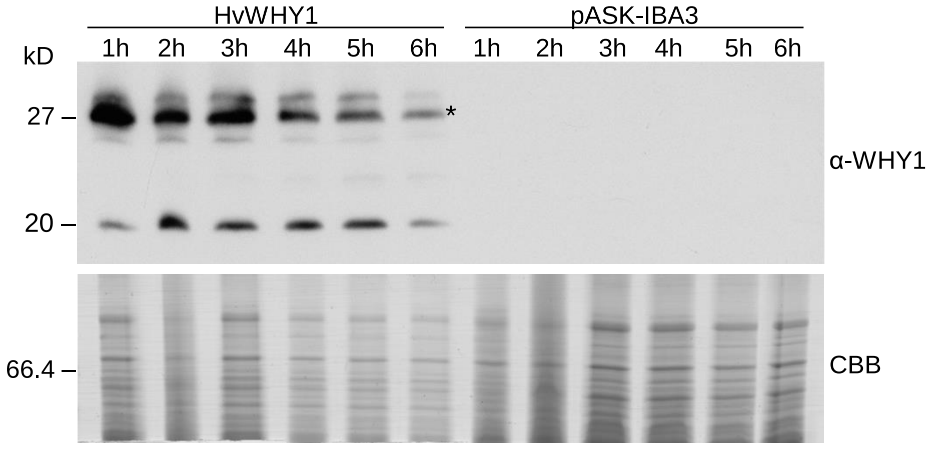


**Figure S1.** Immunological analysis of bacterial cell lysates expressing the coding sequence of *HvWHIRLY1* (HvWHY1) one to six hours (1h-6h) after induction of overexpression using an antibody directed against the HvWHIRLY1 protein. As control lysates of cells containing the empty vector pASK-IBA3 one to six hours (1h-6h) after induction were used. The band marked with an asterisk represents the recombinant HvWHIRLY1 protein. The other bands represent degraded or modified forms of the recombinant HvWHIRLY1. The upper part of the SDS-PA gel was used for staining with Coomassie Brilliant Blue (CBB).

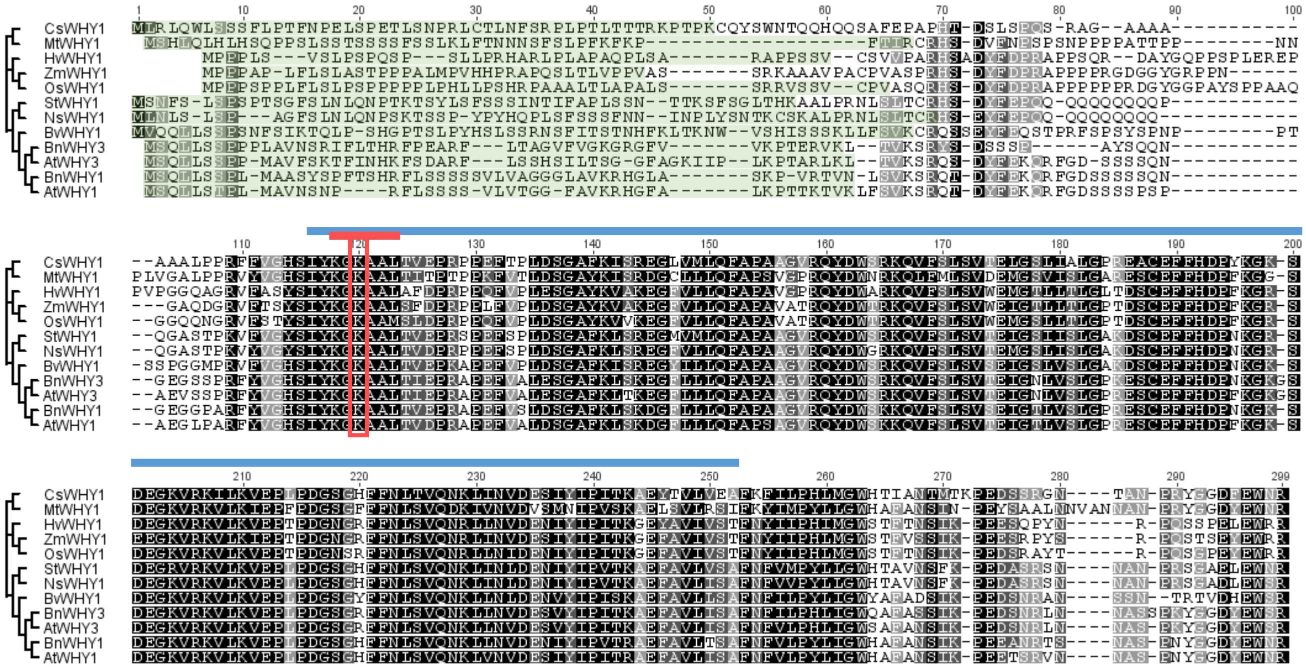


**Figure S2.** Sequence alignment with integrated unrooted neighbor-joining tree of WHIRLY1 and WHIRLY3 proteins. Conserved amino acids are shown in black (100% identity), dark grey (80-100% similarity) and light grey (60-80% similarity), respectively. The following protein sequences were used for the alignment (name, organism, GenBank accession number): CsWHY1, *Cucumis sativus*, XP_004167189; MtWHY1, *Medicago truncatula*, KEH19793; HvWHY1, *Hordeum vulgare*, BAJ96655; ZmWHY1, *Zea mays*, NP_001123589; OsWHY1, *Oryza sativa*, BAD68418; StWHY1, *Solanum tuberosum*, NP_001275155; NsWHY1, *Nicotiana sylvestris*, XP_009758353; BvWHY1, *Beta vulgaris*, XP_010683246; BnWHY3, *Brassica napus*, CDY45200; AtWHY3, *Arabidopsis thaliana*, NP_178377; BnWHY1, *Brassica napus*, CDY66532; AtWHY1, *Arabidopsis thaliana*, NP_172893. The plastid target peptide predicted with TargetP is highlighted in green, the WHIRLY domain is marked with a blue bar, the DNA binding motif (Lys-Gly-Lys-Ala-Ala-Leu) is highlighted in red and the second lysine within the DNA binding motif is framed in red.

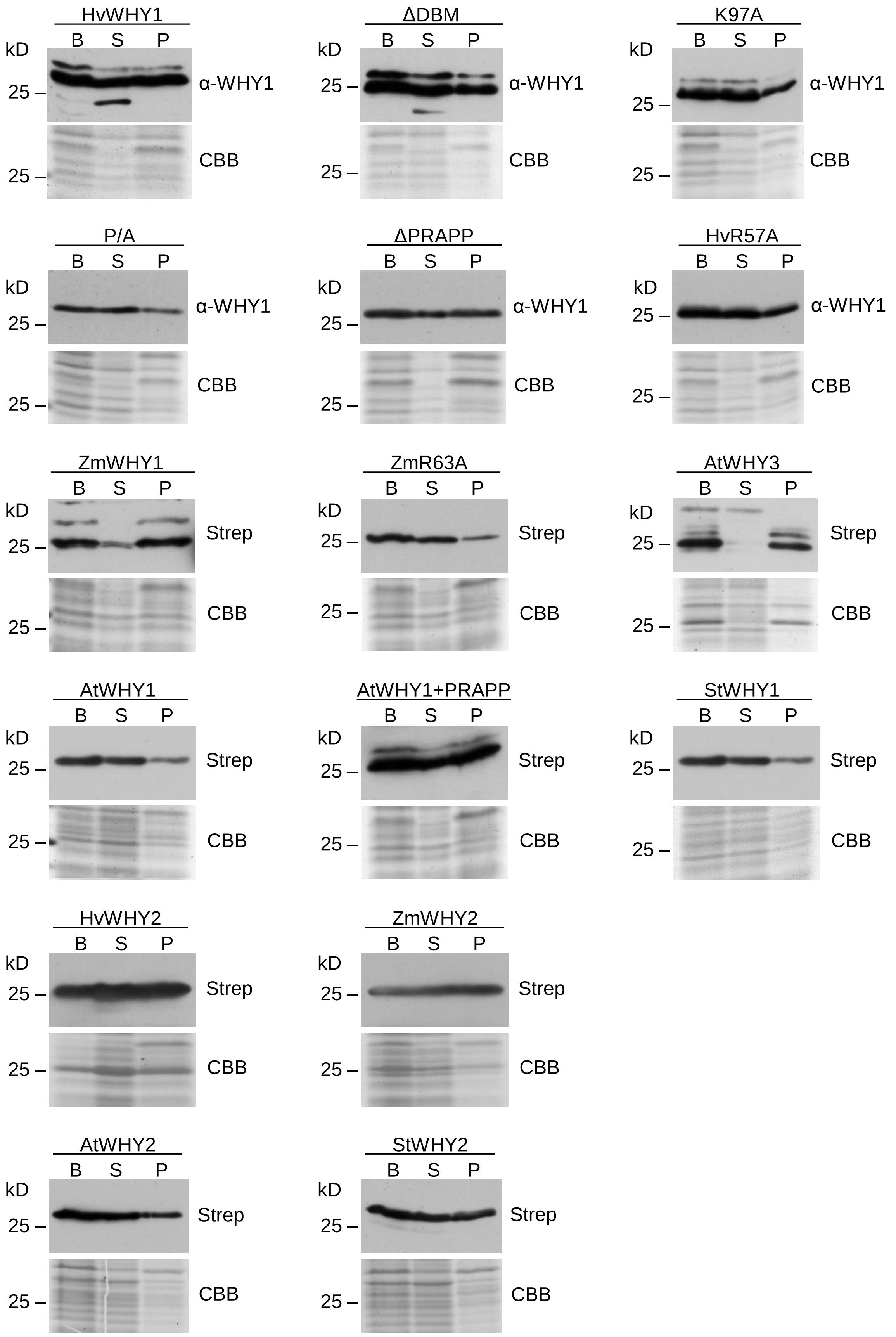


**Figure S3.** Immunological analyses of recombinant protein solubility. Two hours after induction of overexpression bacterial cells (B) were sonicated and afterwards centrifuged to separate the soluble protein fraction (supernatant after sonication and centrifugation (S)) from the unsoluble protein fraction (pellet after sonication and centrifugation (P)). Recombinant proteins were detected with an antibody directed against the HvWHIRLY1 protein (α-WHY1) or with Strep-Tactin (Strep) binding to the C-terminal Strep tag. Comparable gels were stained mit Coomassie Brilliant Blue (CBB).

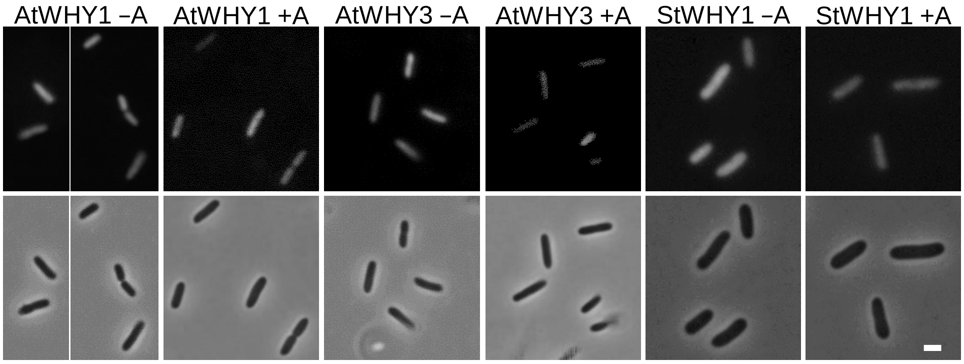


**Figure S4.** Visualization of bacterial nucleoids after accumulation of AtWHIRLY1, AtWHIRLY3 and StWHIRLY1. *E. coli* DH5α cells were transformed either with the pASK-IBA3 vector containing the gene encoding the mature WHIRLY1 or WHIRLY3 protein of *Arabidopsis thaliana* (AtWHY1, AtWHY3) and the gene encoding the mature WHIRLY1 protein of *Solanum tuberosum* (StWHY1). Nucleoids were visualized using DAPI two hours after induction (upper row). Phase contrast images are shown in the lower row. Bar = 2 µm. −A = no anhydrotetracycline, +A = 200 µg/l anhydrotetracycline.

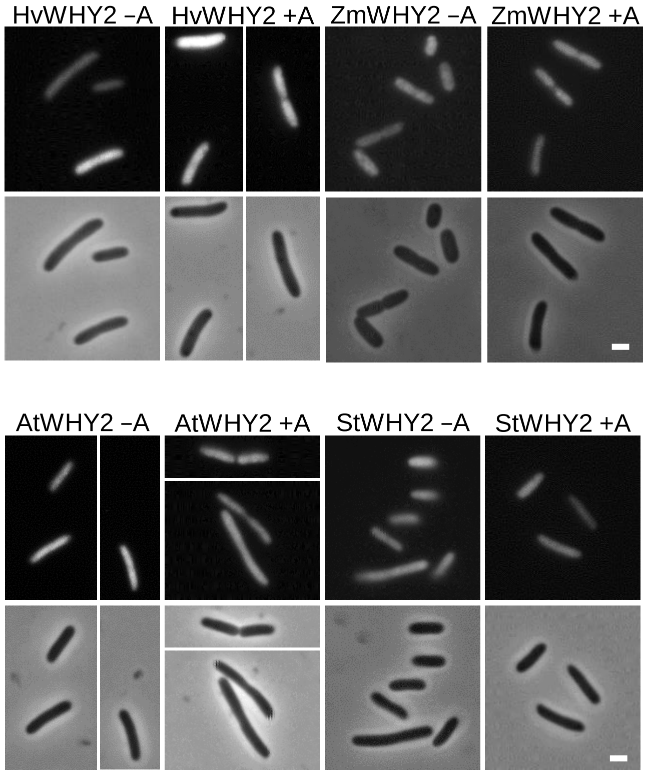


**Figure S5.** Visualization of bacterial nucleoids after accumulation of different WHIRLY2 proteins. *E. coli* DH5α cells were transformed either with the pASK-IBA3 vector containing the gene encoding the mature WHIRLY2 protein of *Hordeum vulgare* (HvWHY2), *Zea mays* (ZmWHY2), *Arabidopsis thaliana* (AtWHY2) or *Solanum tuberosum* (StWHY2). Nucleoids were visualized using DAPI two hours after induction (upper row). Phase contrast images are shown in the lower row. Bar = 2 µm. −A = no anhydrotetracycline, +A = 200 µg/l anhydrotetracycline.

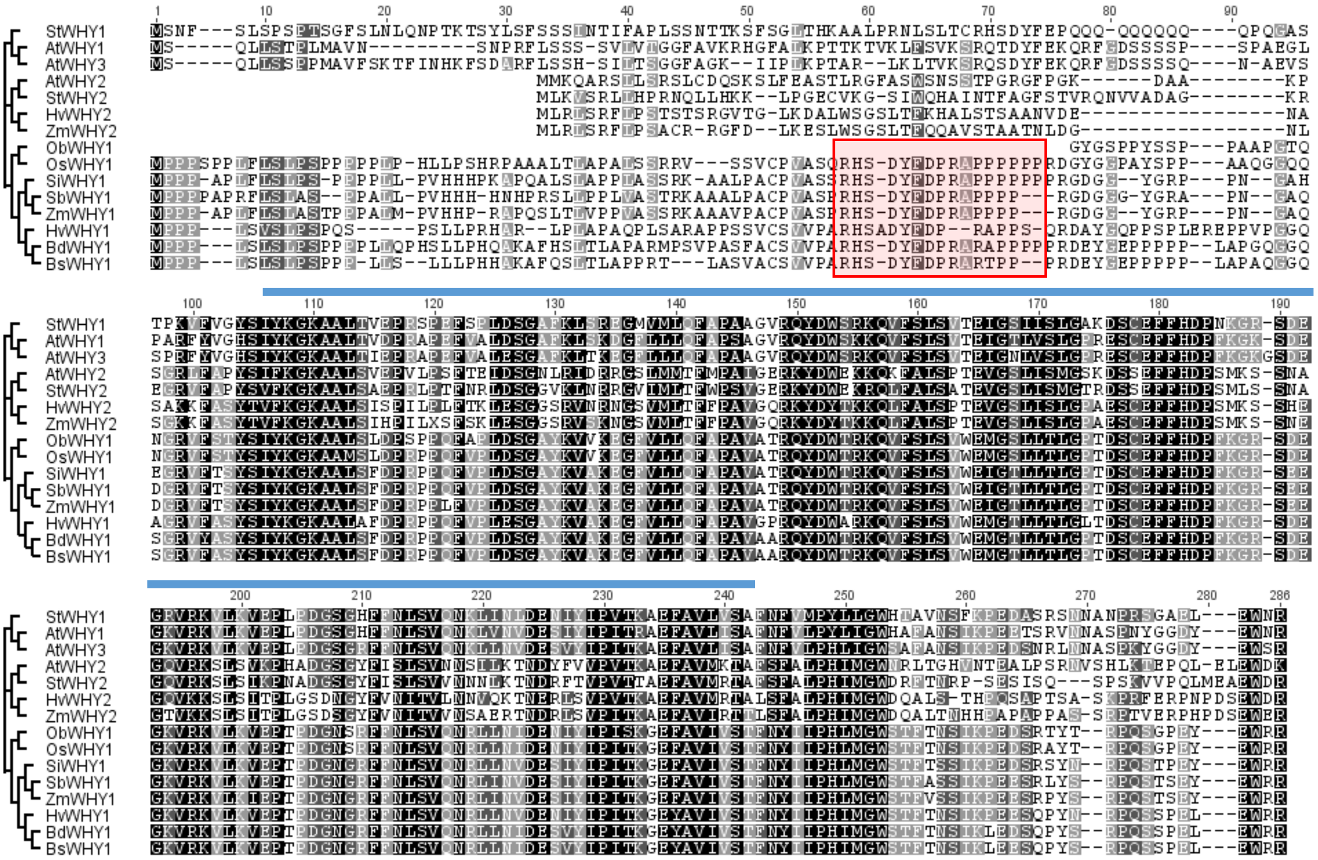

**Figure S6.** Sequence alignment with integrated unrooted neighbor-joining tree of different WHIRLY proteins. Conserved amino acids are shown in black (100% identity), dark grey (80-100% similarity) and light grey (60-80% similarity), respectively. The following protein sequences were used for the alignment (name, organism, GenBank accession number): StWHY1, *Solanum tuberosum*, NP_001275155; AtWHY1, *Arabidopsis thaliana*, NP_172893; AtWHY3, *Arabidopsis thaliana*, NP_178377; AtWHY2, *Arabidopsis thaliana*, NP_177282.2; StWHY2, *Solanum tuberosum*, NP_001275393.1; HvWHY2, *Hordeum vulgare*, BF627441.2; ZmWHY2, *Zea mays*, NP_001152589; ObWHY1, *Oryza brachyantha*, XP_006656631; OsWHY1, *Oryza sativa*, BAD68418; SiWHY1, *Setaria italica*, XP_004964537; SbWHY1, *Sorghum bicolor*, XP_002436467; ZmWHY1, *Zea mays*, NP_001123589; HvWHY1, *Hordeum vulgare*, BAJ96655; BdWHY1, *Brachypodium distachyon*, XP_003557198; BsWHY1, *Brachypodium sylvaticum*, ABL85062. The conserved N-temrinal region in WHIRLY1 proteins of poaceae is highlighted in red and the WHIRLY domain is marked with a blue bar.

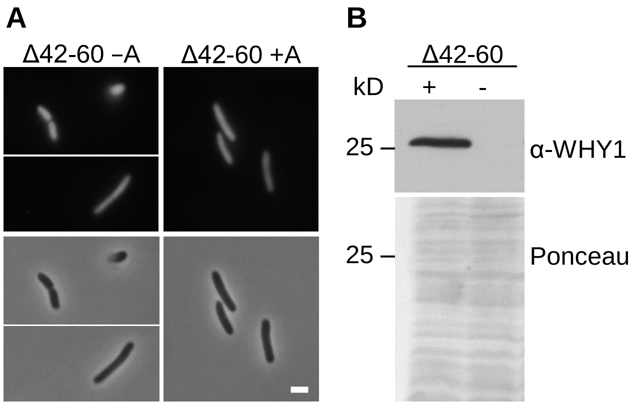


**Figure S7.** Visualization of bacterial nucleoids after accumulation of HvWHIRLY1Δ42-60 and immunological analysis of bacterial cell lysates. *E. coli* DH5α cells were transformed with the pASK-IBA3 vector containing the gene encoding the mutated HvWHIRLY1Δ42-60 protein (Δ42-60). (A) Nucleoids were visualized using DAPI two hours after induction (upper row). Phase contrast images are shown in the lower row. Bar = 2 µm. −A = no anhydrotetracycline, +A = 200 µg/l anhydrotetracycline. (B) Immunological analysis of bacterial cell lysates expressing the coding sequence of *HvWHIRLY1Δ42-60* for two hours (+) or of lysates taken from cells without inducing of overexpression collected at the same time (−) using an antibody directed against the HvWHIRLY1 protein. Prior to immunological detection the membrane was stained with Ponceau.

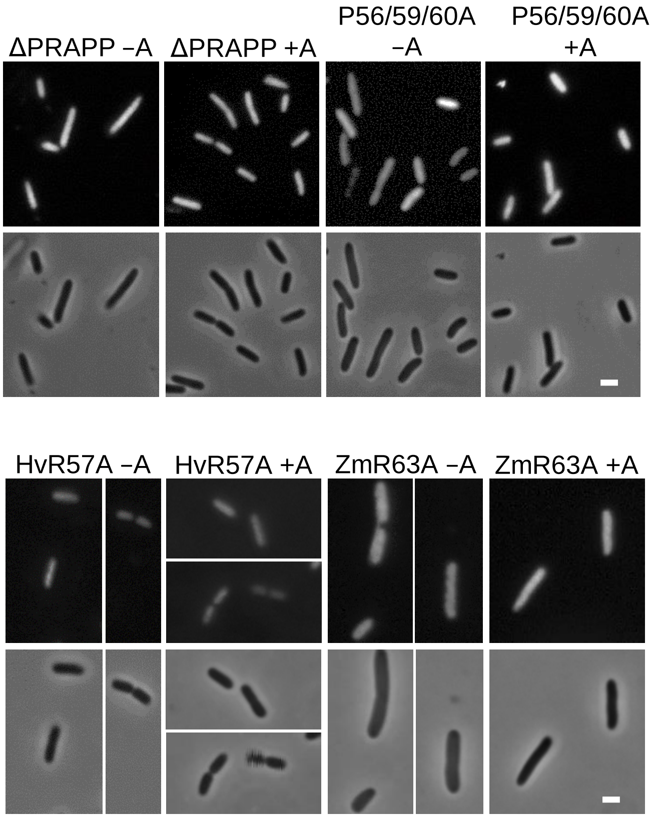


**Figure S8.** Visualization of bacterial nucleoids after accumulation of HvWHIRLY1 proteins with altered PRAPP motifs. *E. coli* DH5α cells were transformed either with the pASK-IBA3 vector containing the gene encoding the HvWHIRLY1ΔPRAPP protein (ΔPRAPP), the HvWHIRLY1P56/59/60A protein (P56/59/60A), the HvWHIRLY1R57A protein (HvR57A) or ZmWHIRLY1R63A protein (ZmR63A). Nucleoids were visualized using DAPI two hours after induction (upper row). Phase contrast images are shown in the lower row. Bar = 2 µm. −A = no anhydrotetracycline, +A = 200 µg/l anhydrotetracycline.

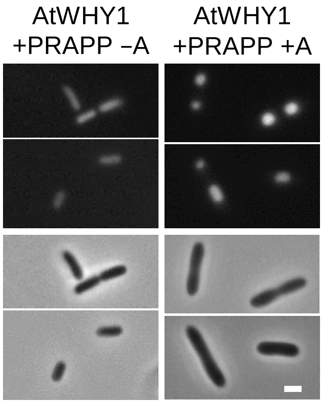


**Figure S9.** Visualization of bacterial nucleoids after accumulation of the chimeric AtWHIRLY1+PRAPP protein. *E. coli* DH5α cells were transformed with the pASK-IBA3 vector containing the gene encoding the AtWHIRLY1+PRAPP protein (AtWHY1+PRAPP). Nucleoids were visualized using DAPI two hours after induction (upper row). Phase contrast images are shown in the lower row. Bar = 2 µm. −A = no anhydrotetracycline, +A = 200 µg/l anhydrotetracycline.

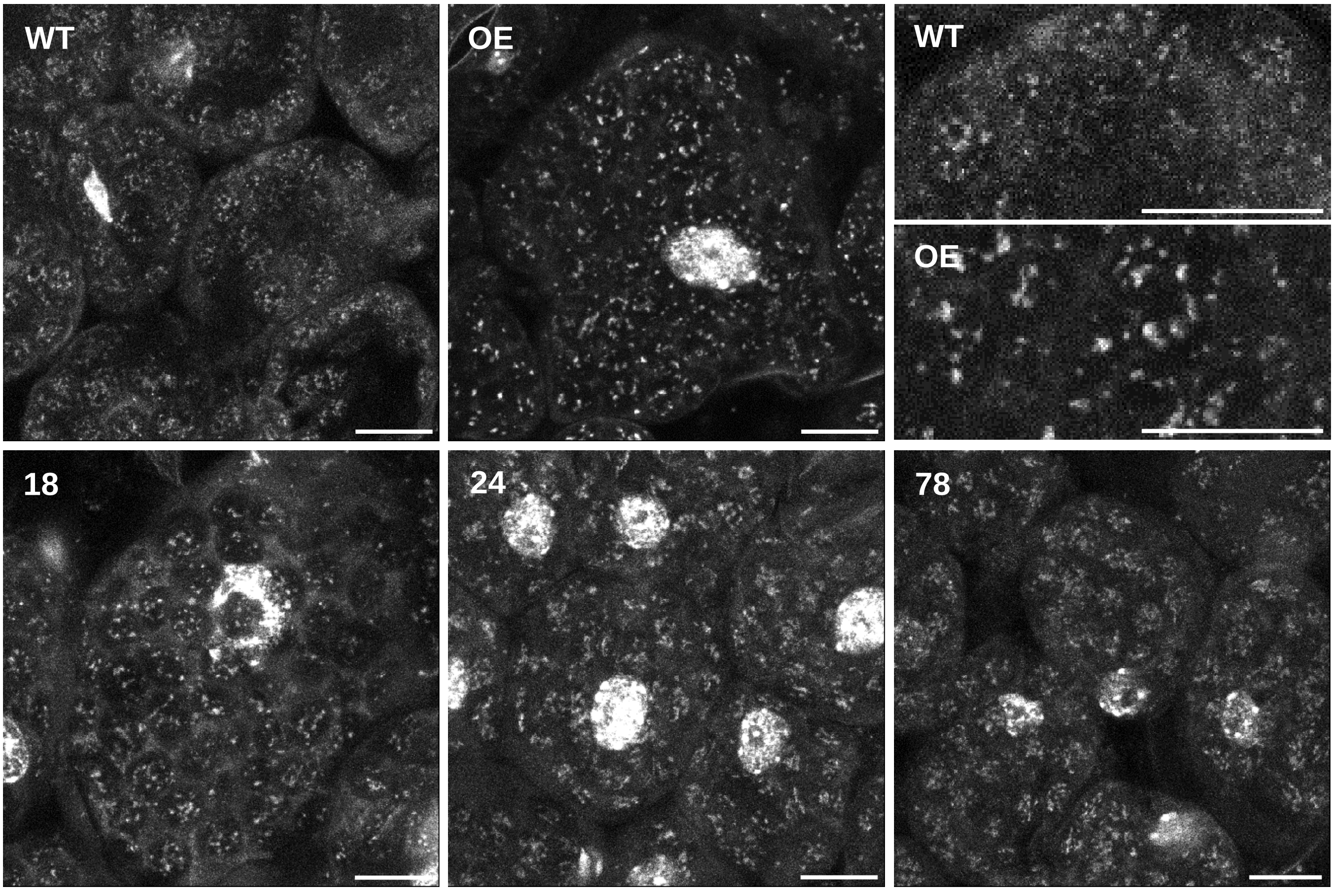


**Figure S10.** Comparison of the nucleoid morphology in different *WHIRLY1* overexpression lines. Nucleoid morphology in leaves of the wild type (WT), the AtWHIRLY1:HA overexpression line (OE) and the AtWHIRLY1+PRAPP:HA overexpression lines 18, 24 and 78 is shown. Staining of DNA was performed with SYBR Green I on cross-sections prepared from the first true leaves of seedlings 14 days after sowing. Microscopy was performed with a confocal laser scanning microscope. Bars = 10 µm.

### Supplementary Tables

**Table S1.** Gene specific primers that were used for amplification of *WHIRLY* coding sequences encoding the mature protein for generation of *E. coli* overexpression constructs. *Truncated *HvWHIRLY1* coding sequence at the 5'-end from nucleotide 124-180 resulting in deletion of amino acids 42-60.

| Gene name | Accession no. | Primer name | Sequence |
|---|---|---|---|
| *HvWHIRLY1* | AK365452.1 | HvWHY1-pASK3_for | 5'-atggtaggtctcaaatgatgtgctccgtcgtccccgc-3' |
| | | HvWHY1-pASK3_rev | 5'-atggtaggtctcagcgctccttcgccactcaattcggg-3' |
| *HvWHIRLY1* (Δ42-60)* | AK365452.1 | HvWHY1-del124-180_for | 5'-atggtaggtctcaaatgatgtcgcagcgcgacgc-3' |
| | | HvWHY1-pASK3_rev | 5'-atggtaggtctcagcgctccttcgccactcaattcggg-3' |
| *HvWHIRLY2* | BF627441.2 | pASK-3_HvWHY2_for | 5'-atggtaggtctcaaatgatggatgagaatgcatctgct aag-3' |
| | | pASK-3_HvWHY2_rev | 5'-atggtaggtctcagcgctcctgtcccattcagaatctg ggt-3' |
| *ZmWHIRLY1* | Zm.88681 | ZmWHY1-pASK3_for | 5'-atggtaggtctcaaatggcctcctcccgtaaggccgc-3' |
| | | ZmWHY1-pASK3_rev2 | 5'-atggtaggtctcagcgctccgacgccattcgtactca gac-3' |
| *ZmWHIRLY2* | NM_001159117.2 | ZmWHY2-pASK_for | 5'-atggtaggtctcaaatgctgcggctctcccgcttc-3' |
| | | ZmWHY2-pASK_rev | 5'-atggtaggtctcagcgctcctctcccattcggaatcc-3' |
| *AtWHIRLY1* | At1g14410 | AtWHY1-pASK-IBA3_for | 5'-atggtaggtctcaaatgatggtgaagctcttctcggt-3 |
| | | AtWHY1-pASK-IBA3_rev | 5'-atggtaggtctcagcgcttctattccattcatagtctcct-3 |
| *AtWHIRLY2* | At1g71260 | pASK-3_AtWHY2_for | 5'-atggtaggtctcaaatgatggcaagctggtcaaattct tca-3 |
| | | pASK-3_AtWHY2_rev | 5'-atggtaggtctcagcgcttttatcccactccagctctaa ctg-3 |
| *AtWHIRLY3* | At2g02740 | AtWHY3-pASK-IBA3_for | 5'-atggtaggtctcaaatgatgttgaaattaacggtga agt-3 |
| | | AtWHY3-pASK-IBA3_rev | 5'-atggtaggtctcagcgcttctactccattcgtagtctc-3 |
| *StWHIRLY1* | NM_001288226.1 | StWHY1-pASK_for | 5'-atggtaggtctcaaatgtccaatttctctctttctccttc-3' |
| | | StWHY1-pASK_rev | 5'-atggtaggtctcagcgcttcgattccattcaagttca gca-3' |
| *StWHIRLY2* | NM_001288464.1 | StWHY2-pASK_for | 5'-atggtaggtctcaaatgttgaaagtcagtcggcttctg-3' |
| | | StWHY2-pASK_rev | 5'-atggtaggtctcagcgcttctatcccactccgcctcc-3' |

**Table S2.** Primers that were used for site-directed mutagenesis.

| Gene name | Mutation | Primer name | Sequence |
|---|---|---|---|
| *HvWHIRLY1* (AK365452.1) | Deletion of DNA binding motif (KGKAAL) | HvWHY1ΔDBM-ivM_for | 5'-agcatctacaagggcgccgccgcgctggccttc-3' |
| | | HvWHY1ΔDBM-ivM_rev | 5'-gaaggccagcgcggcggcgcccttgtagatgct-3' |
| | Substitution of lysine 97 with an alanine | HvWHY1K97A-ivM-for | 5'-agctacagcatctacgccttcgaccccagg-3' |
| | | HvWHY1K97A-ivM-rev | 5'-cctggggtcgaaggcgtagatgctgtagct-3' |
| | Deletion of the PRAPP motif | Δ56-60_rev | 5'-gcgtcgcgctgcgagtcgaagtagtcgg-3' |
| | | Δ56-60_for | 5'-ccgactacttcgactcgcagcgcgacgc-3' |
| | Substitution of proline 56, 59 and 60 with an alanine | P56/59/60A_for | 5'-acttcgacgcccgggccgcggcgtcgcag-3' |
| | | P56/59/60A_rev | 5'-ctgcgacgccgcggcccgggcgtcgaagt-3' |
| | Substitution of arginine 57 with an alanine | ivM_HvWHY1R57A_for | 5'-tacttcgaccccgcggccccgccgtc-3' |
| | | ivM_HvWHY1R57A_rev | 5'-gacggcggggccgcggggtcgaagta-3' |
| *ZmWHIRLY1* (Zm.88681) | Substitution of arginine 63 with an alanine | ivM_ZmWHY1R63A_for | 5'-ctacttcgatccggcggctccgccgccg-3' |
| | | ivM_ZmWHY1R63A_rev | 5'-cggcggcggagccgccggatcgaagtag-3' |
| *AtWHIRLY1* (At1g14410) | Insertion of the PRAPP motif | AtWHY1+PRAPP_for | 5'-gacggattacttcgagccccgggccccgccgaagcagaggttcggtg-3' |
| | | AtWHY1+PRAPP_rev | 5'-caccgaacctctgcttcggcggggcccggggctcgaagtaatccgtc-3' |

**Table S3.** Gene specific primers that were used for qRT-PCR analysis of transcript levels in *E. coli*.

| Gene name | Accession no. | Primer name | Sequence |
|---|---|---|---|
| *cysG* | 947880 | Ec-cysG_for | 5‘-gaaaggtggcgatccgttta-3‘ |
| | | Ec-cysG_rev | 5‘-atacccgaataggcagagca-3‘ |
| *dnaA* | 948217 | Ec-dnaA_for | 5‘-aaagtcgcggatctcctttc-3‘ |
| | | Ec-dnaA_rev | 5‘-ggcagactgtggttagtcag-3‘ |
| *ftsZ* | 944786 | Ec-ftsZ_for | 5‘-cggtatcaccaaaggactgg-3 |
| | | Ec-ftsZ_rev | 5‘-tacccgcagcaataaagacc-3 |
| *hns* | 945829 | Ec-hns_for | 5‘-tattgacccgaacgaactgc-3‘ |
| | | Ec-hns_rev | 5‘-tagtttcgccgttttcgtca-3‘ |
| *hupA* | 948499 | Ec-hupA_for | 5’-ctctgaaagaaggcgatgct-3’ |
| | | Ec-hupA_rev | 5’-cagaaacaaatgccggtacg-3’ |
| *idnT* | 948798 | Ec-idnT_for | 5‘-tgatttctgatacgggtgcg-3‘ |
| | | Ec-idnT_rev | 5‘-agcaggacaaaacccacttc-3‘ |
| *rpoA* | 947794 | Ec-rpoA_for | 5‘-gtggagcgtattgcctacaa-3‘ |
| | | Ec-rpoA_rev | 5‘-gaatcgcctcttcaggatcg-3‘ |
